# Structural and molecular basis of resistance to aminoglycoside antibiotics by APH(3’)-IIb from *Pseudomonas aeruginosa*

**DOI:** 10.1101/2025.03.30.646141

**Authors:** Julien Kowalewski, Mathilde Tomaszczyk, Jean-François Guichou, Muriel Gelin, Gilles Labesse, Corinne Lionne

## Abstract

*Pseudomonas aeruginosa* is one of the six bacteria of greatest concern identified by the WHO for its resistance to antibiotics. In the case of aminoglycosides, used in combination with other antibiotics in the treatment of severe infections, resistance is mainly due to modification of the antibiotic by bacterial enzymes. In this study, we have functionally and structurally characterized one such enzyme from *P. aeruginosa*, aminoglycoside 3’-*O*-phosphotransferase IIb, APH(3’)-IIb. The results will provide a better understanding of this resistance mechanism and enable us to envisage solutions for blocking it and restoring the efficacy of aminoglycosides.

## Introduction

*Pseudomonas aeruginosa* is a Gram-negative opportunistic bacterium, generally harmless in healthy people, but capable of causing serious infections in the weakened. *P. aeruginosa* is a frequent cause of nosocomial infections such as healthcare- or ventilator-associated pneumonia, bacteremia, urinary tract infections or surgical or burn wounds [1]. Patients with cystic fibrosis are particularly prone to pyocyanic lung infections. The pulmonary inflammation associated with bacterial infection leads to severe damage that can necessitate transplantation and, in the most severe cases, death [2,3]. The recommended treatment for these infections is a combination of two antibiotics, usually a β-lactam and an aminoglycoside [1,4,5].

Unfortunately, as highlighted by the WHO since 2017, *P. aeruginosa* is among the critical human pathogens in terms of resistance. In fact, it is one of the 6 antibiotic-resistant “priority pathogens” for which new antibiotics are urgently required [6,7]. *P. aeruginosa* exhibits a high level of intrinsic resistance involving several mechanisms [8,9]: restricted outer membrane permeability, efflux pumps and antibiotic-inactivating enzymes, β-lactamases and aminoglycoside-modifying enzymes (AMEs). In addition, it has also acquired antibiotic resistance though mutations and horizontal transfer of antibiotic resistance genes, including AMEs. Its ability to form biofilms in which persisters are protected from antimicrobial agents gives it additional adaptive resistance to antibiotics [10]. Even one of the most recently discovered antibiotics effective against human pathogens, clovibactin, is ineffective against *P. aeruginosa* [11].

Aminoglycosides are broad-spectrum antibiotics whose use is mainly reserved for the treatment of severe infections [12]. This is particularly the case for cystic fibrosis patients suffering from *P. aeruginosa* infections, who are treated with inhaled amikacin or tobramycin [13]. As with other antibiotics, the development of resistance to aminoglycosides makes it difficult to treat the infection [14]. Various resistance mechanisms exist in this bacterium, including modification of the aminoglycoside by enzymes [15,16]. These enzymes are classified into three families according to the modification they bring to the aminoglycoside. Aminoglycoside acetyltransferases (AAC) use acetyl coenzyme A to transfer an acetyl group to an amine of the aminoglycoside. Aminoglycoside adenylyltransferases (ANT) and phosphotransferases (APH) use a nucleotide to transfer a nucleotidyl residue or a phosphate respectively to a hydroxyl of the aminoglycoside [16,17]. These modifications render the antibiotic incapable of interacting with its target, bacterial 16S RNA.

In this study, we characterized an APH present in *P. aeruginosa* that catalyzes phosphate transfer onto the 3’-hydroxyl of aminoglycoside, APH(3’)-IIb. This enzyme is widely present in clinical isolates of *P. aeruginosa*, particularly in burn victims and cystic fibrosis patients: 45% and 90% respectively [18]. There is no human orthologues of APHs and, despite the structural similarity of their nucleotide-binding site to that of eukaryotic kinases, APH inhibitors could be considered as resistance-reversing agents [19–24]. Thus, a better understanding of the functional and structural properties of APH(3’)-IIb would help guide the design of effective inhibitors.

The *aph(3’)-IIb* gene was first described in 1996 and suggested to have a chromosomal localization [25]. The presence of the gene in the chromosome of the strain PA01 was confirmed during complete genome sequencing [26]. Because it was found in all *P aeruginosa* isolates tested, it was considered as responsible for this bacterium “uniform resistance” to kanamycin. Later, it was shown that *aph(3’)-IIb* gene is under the positive control of the HpaA surrogate regulator, suggesting a correlation between aminoglycoside resistance and carbon metabolism [27]. Indeed, *P aeruginosa* can use 4-hydroxyphenylacetic acid (HPA) as its sole carbon source and, via an *hpa* pathway, it can also use dopamine and tyramine, which are aromatic compounds present in the mammalian nervous system. The same study showed that coexistence of *hpaA* and *aph* genes is a common feature in *P. aeruginosa,* and that a higher proportion of isolates from the clinic than from the environment showed increased HPA-induced resistance to neomycin. This link between antibiotic resistance and metabolism may be particularly important for the environmental adaptation of bacteria in the context of a human infection.

Two previous biochemical characterizations of purified APH(3’)-IIb showed specificity for kanamycin, neomycin, paromomycin, ribostamycin and butirosin [28,29]. One proposed explanation for the lack of phosphorylation of amikacin by APH(3’)-IIb is that the presence of the L-hydroxyaminobuteroyl amide (L-HABA) substitution at N1 may prevent interactions in the active site. Furthermore, APH(3’)-IIb can only phosphorylate neomycin on the 3’-hydroxyl, unlike other members of the APH(3’) sub-family, which can also catalyze 5’’-phosphorylation of this aminoglycoside. From a structural point of view, little is known about APH(3’)-IIb, except that unlike APH(3’)-Ia and -IIIa, which form dimer in solution in the absence of reducing agents [30,31], APH(3’)-IIb has been shown to be monomeric. Here, we investigated the substrate specificity of APH(3’)-IIb, not only for aminoglycosides, but also for the phosphate donor. We confirmed that it is monomeric in solution and determined the crystal structure of two variants, the wild-type and a spontaneous single mutant of the gatekeeper residue. We obtained a structure for the APH(3’)-IIb·MgADP·kanamycin-phosphate ternary complex after crystals of the apo protein were soaked in MgATP and kanamycin and catalysis took place in the crystal.

The results of this study will form the basis for the design of specific APH inhibitors that could be used as adjuvants to aminoglycoside therapy, in the same way that β-lactamase inhibitors are used in combination with β-lactam antibiotics [32–34].

## Results

When cloning *aph(3’)-IIb* into the expression vector, we realized that a spontaneous point mutation of the gatekeeper residue had occurred, transforming methionine 95 into leucine. Among the various APH subfamilies, the nature of the gatekeeper residue has been shown to influence the specificity of the enzyme for the phosphate donor substrate [35–37]. Thus, APHs containing a methionine gatekeeper residue are specific for ATP, those with a tyrosine are specific for GTP and those with a phenylalanine can use either nucleotide. It has been suggested that the bulky side chain of tyrosine is too close to the amine at position 6 of the adenine ring, blocking access to the binding pocket for ATP. Here, we compare the functional and structural properties of the two variants of the APH(3’)-IIb, carrying either a methionine or a leucine as gatekeeper residue, and therefore both predicted to be specific for ATP.

### The APH(3’)-IIb WT and M95L variants are specific for MgATP as a phosphate donor

First, we quantified the enzymatic activity of the two variants using ATP or GTP as the phosphate donor and kanamycin A as the acceptor. Phosphorylation of kanamycin A is more efficient with MgATP than with MgGTP as phosphate donor: 96-fold with wild-type APH(3’)-IIb (WT) and 46-fold with the M95L variant (**Fig 1**). The difference in catalytic efficiency is mainly due to a larger K_m_ value for MgGTP than for MgATP, suggesting that the active site of APH(3’)-IIb is not suitable for guanine nucleobase binding. Catalytic efficiencies using MgATP or MgGTP are similar for each variant. However, k_cat_ values are 2.2 to 3.5-fold higher with the M95L variant than with the WT, but this is compensated by 2.4 to 8-fold higher K_m_ values.

**Fig 1.**
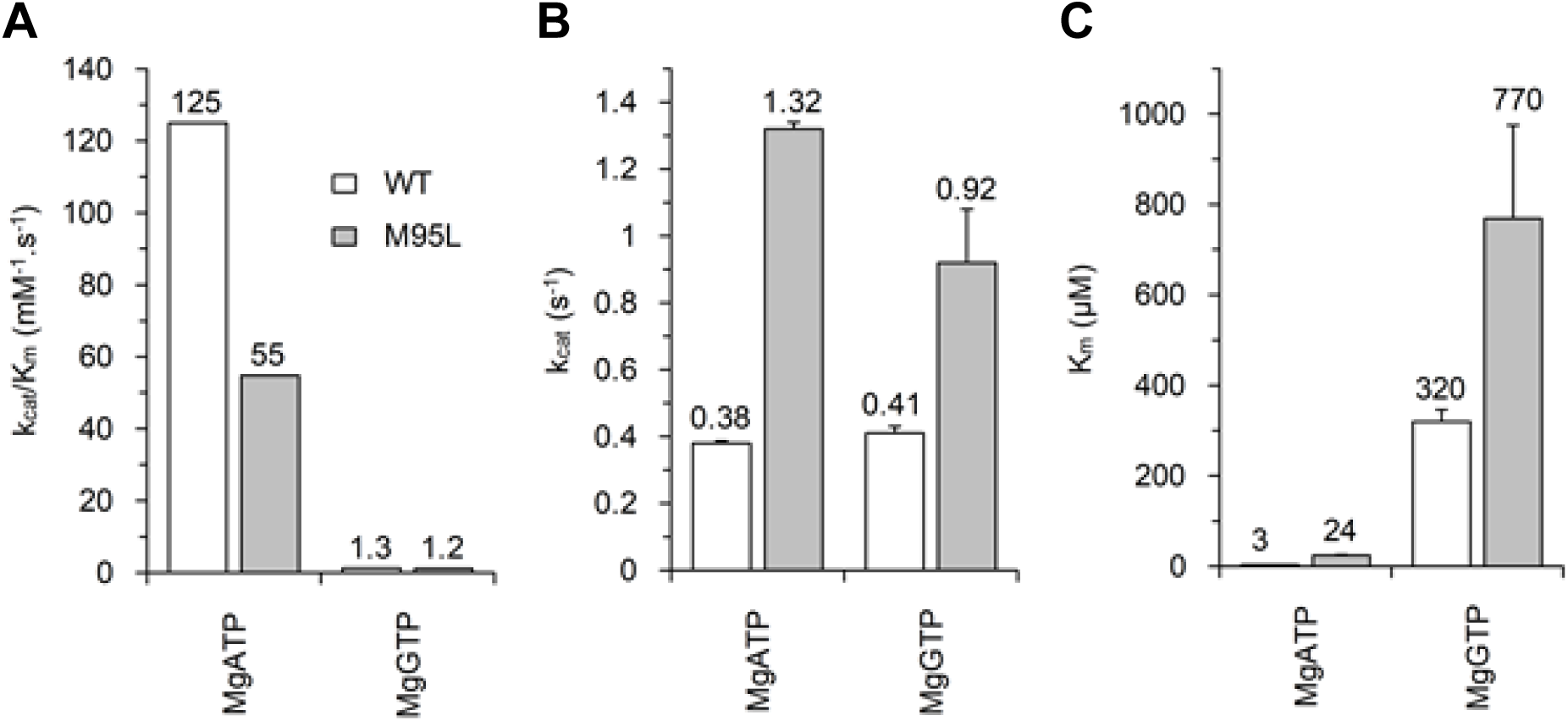
Kinetic parameters for the phosphorylation of kanamycin A by APH(3’)-IIb WT or the M95L variant using MgATP or MgGTP as phosphate donor. (A) Catalytic efficiencies, (B) catalytic constants and (C) Michaelis constants determined at 25°C are shown as white bars for APH(3’)-IIb WT and grey bars for M95L variant. Parameter values are given above the bars.

Next, we determined the affinity of APH(3’)-IIb WT or M95L variants for different adenosine- or guanosine-containing nucleotides using ITC (**S1 Fig**). With adenosine-containing nucleotides, the K_d_ values are 1.7 to 2.9-fold larger with the M95L variant than with the WT, but these differences in affinity between the two variants are not found with guanosine-containing nucleotides (**Fig 2A** and **S1 Table**). This again suggests that the active site of APH(3’)-IIb cannot easily accommodate nucleotides with a guanine base. With adenosine-containing nucleotides, the binding is clearly enthalpy-driven (**Fig 2B**), but not with guanosine-containing nucleotides for which both enthalpy and entropy contribute to the interactions.

**Fig 2.**
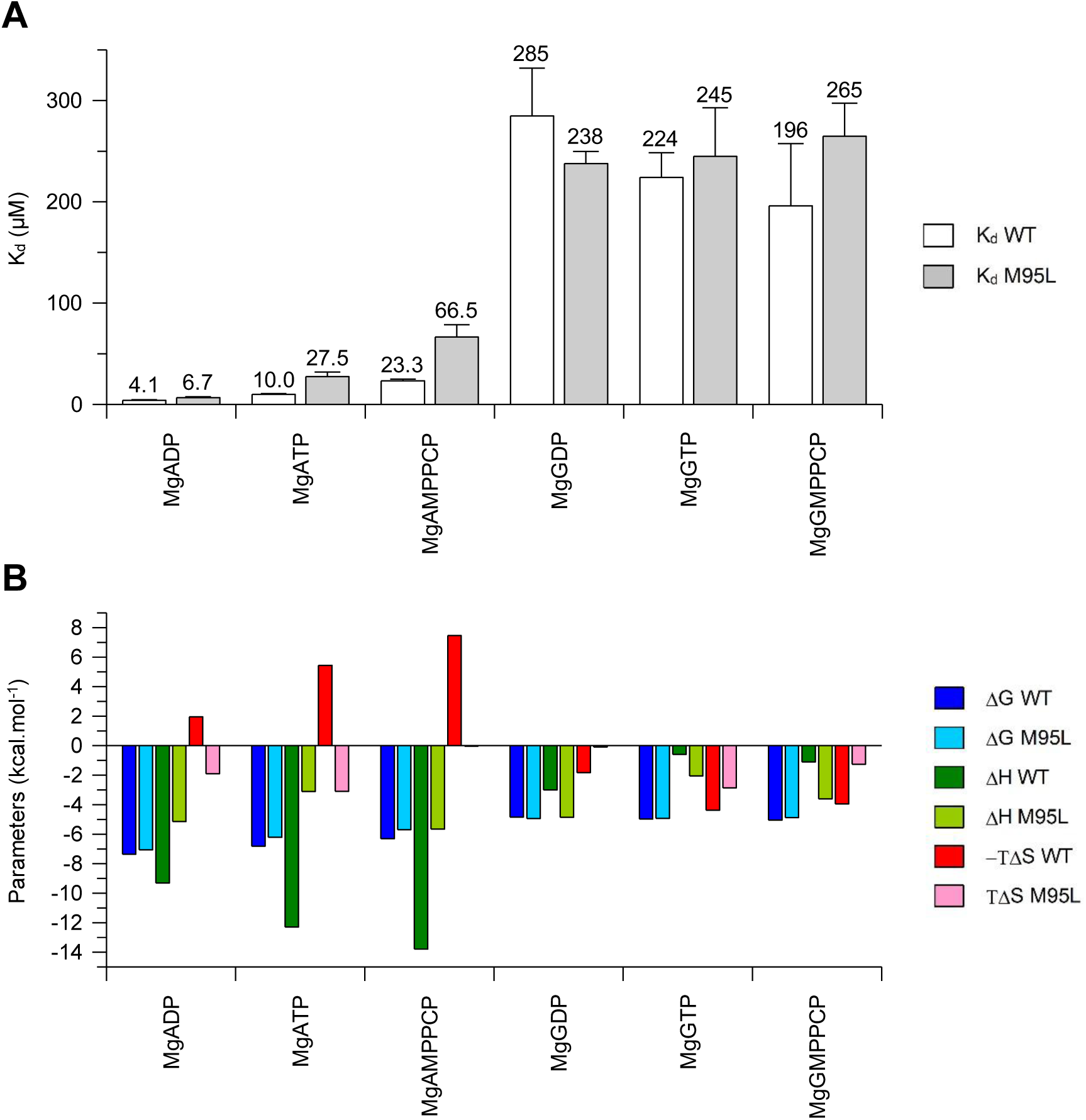
Dissociation constants and thermodynamic parameters for the binding of different nucleotides to APH(3’)-IIb WT or M95L. (**A**) K_d_ for nucleotides with APH(3’)- IIb WT (white bars) and the M95L variant (grey bars). (**B**) Thermodynamic parameters for the binding of nucleotides to APH(3’)-IIb WT (dark colors) or to the M95L variant (light colors). The calculated Gibb’s free energy (ΔG) is shown in blue, the binding enthalpy (ΔH) in green and the binding entropy (–TΔS) in red. Values of K_d_, stoichiometry (N) and thermodynamic parameters are given in **S1 Table** and binding isotherms are shown in **S1 Fig**.

### APH(3’)-IIb has low specificity for aminoglycosides

We then characterized the specificity of APH(3’)-IIb for different aminoglycosides containing 2 or 3 rings on either side of the central 2-deoxystreptamine. The 4,5-disubstituted aminoglycosides containing 4 rings including one ribose (neomycin and paromomycin) are among the most efficiently phosphorylated by the two APH(3’)-IIb variants (**Fig 3** and **Table 1**). This is mainly due to a low K_m_ value. The difference between neomycin and paromomycin lies in the 6’ moiety (highlighted by a blue circle in **Table 1**). The amine carried by neomycin appears to be slightly more favorable than the hydroxyl carried by paromomycin. This is reflected in the differences in affinity measured by ITC (**Fig 4** and **S1 Table**). Ribostamycin, which contains a hydroxyl instead of neomycin’s fourth ring on the 3’ ribose (green box in **Table 1**), has a lower catalytic efficiency, due to a higher K_m_. This correlates with a higher K_d_ measured by ITC.

**Fig 3.**
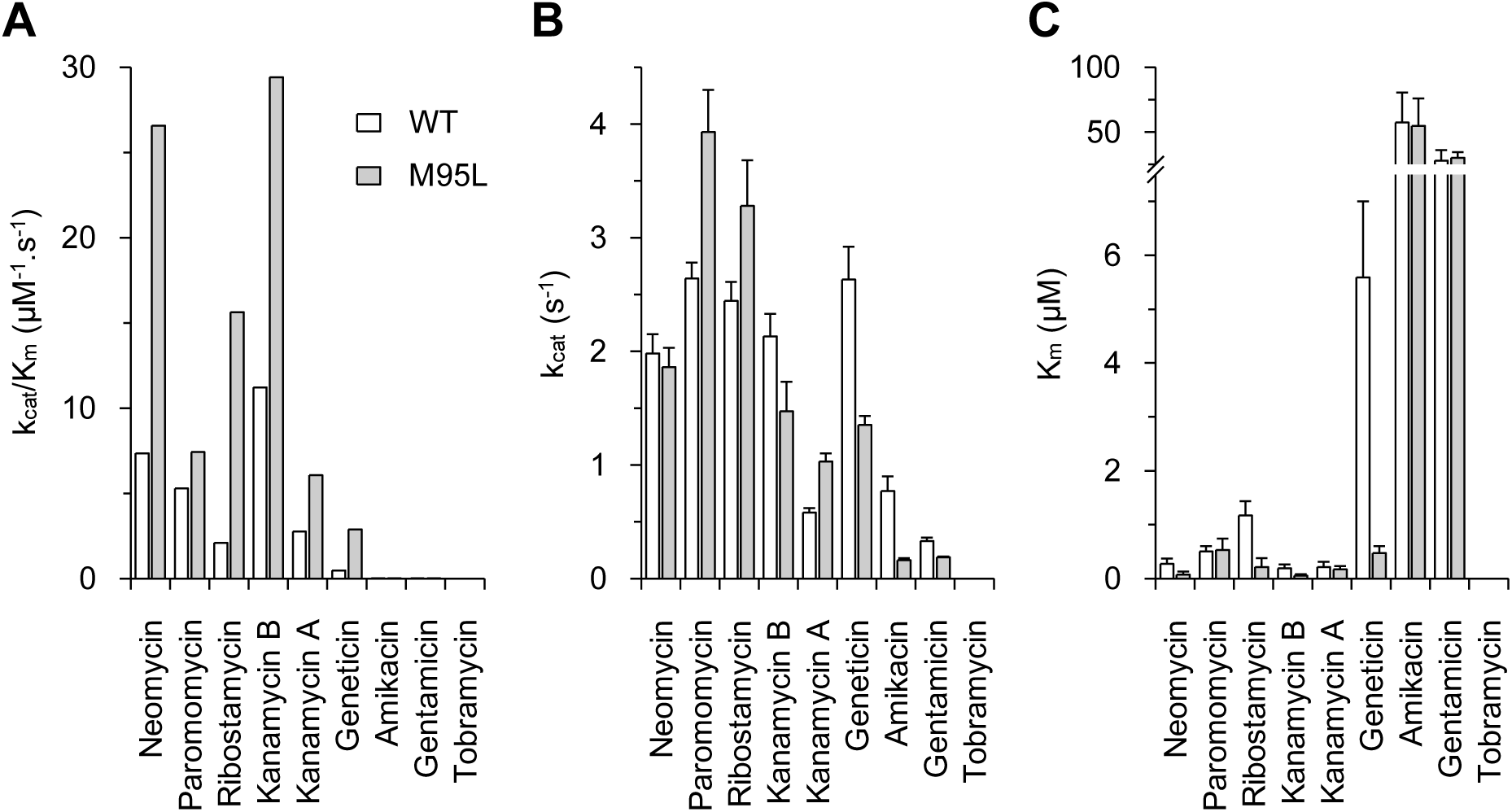
Kinetic parameters for the phosphorylation of different aminoglycosides by APH(3’)-IIb WT or the M95L variant. Values for catalytic constant, k_cat_, Michaelis constant, K_m_, and catalytic efficiency, k_cat_/K_m_, are given in **Table 1**.

**Table 1.**
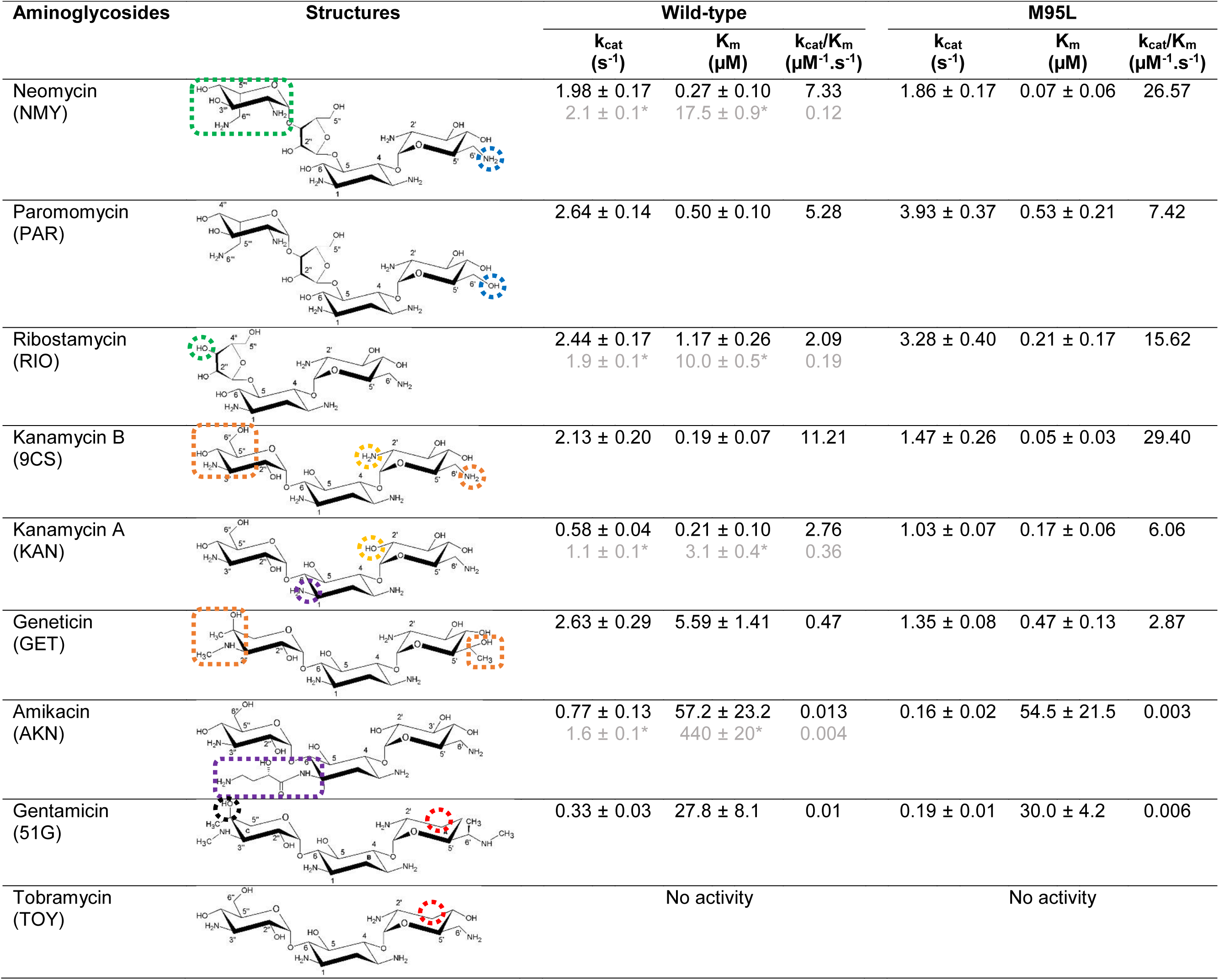
Kinetic parameters for the phosphorylation of different aminoglycosides by APH(3’)-IIb. *Values in grey are from [29].

| Aminoglycosides | Structures | Wild-type |  |  | M95L |  |  |
| --- | --- | --- | --- | --- | --- | --- | --- |
| | | $k_{cat}$<br>(s<sup>-1</sup>) | $K_m$<br>(μM) | $k_{cat}/K_m$<br>(μM<sup>-1</sup>.s<sup>-1</sup>) | $k_{cat}$<br>(s<sup>-1</sup>) | $K_m$<br>(μM) | $k_{cat}/K_m$<br>(μM<sup>-1</sup>.s<sup>-1</sup>) |
| Neomycin<br>(NMY) |  | 1.98 ± 0.17<br>2.1 ± 0.1* | 0.27 ± 0.10<br>17.5 ± 0.9* | 7.33<br>0.12 | 1.86 ± 0.17 | 0.07 ± 0.06 | 26.57 |
| Paromomycin<br>(PAR) |  | 2.64 ± 0.14 | 0.50 ± 0.10 | 5.28 | 3.93 ± 0.37 | 0.53 ± 0.21 | 7.42 |
| Ribostamycin<br>(RIO) |  | 2.44 ± 0.17<br>1.9 ± 0.1* | 1.17 ± 0.26<br>10.0 ± 0.5* | 2.09<br>0.19 | 3.28 ± 0.40 | 0.21 ± 0.17 | 15.62 |
| Kanamycin B<br>(9CS) |  | 2.13 ± 0.20 | 0.19 ± 0.07 | 11.21 | 1.47 ± 0.26 | 0.05 ± 0.03 | 29.40 |
| Kanamycin A<br>(KAN) |  | 0.58 ± 0.04<br>1.1 ± 0.1* | 0.21 ± 0.10<br>3.1 ± 0.4* | 2.76<br>0.36 | 1.03 ± 0.07 | 0.17 ± 0.06 | 6.06 |
| Geneticin<br>(GET) |  | 2.63 ± 0.29 | 5.59 ± 1.41 | 0.47 | 1.35 ± 0.08 | 0.47 ± 0.13 | 2.87 |
| Amikacin<br>(AKN) |  | 0.77 ± 0.13<br>1.6 ± 0.1* | 57.2 ± 23.2<br>440 ± 20* | 0.013<br>0.004 | 0.16 ± 0.02 | 54.5 ± 21.5 | 0.003 |
| Gentamicin<br>(51G) |  | 0.33 ± 0.03 | 27.8 ± 8.1 | 0.01 | 0.19 ± 0.01 | 30.0 ± 4.2 | 0.006 |
| Tobramycin<br>(TOY) |  | No activity |  |  | No activity |  |  |

**Fig 4.**
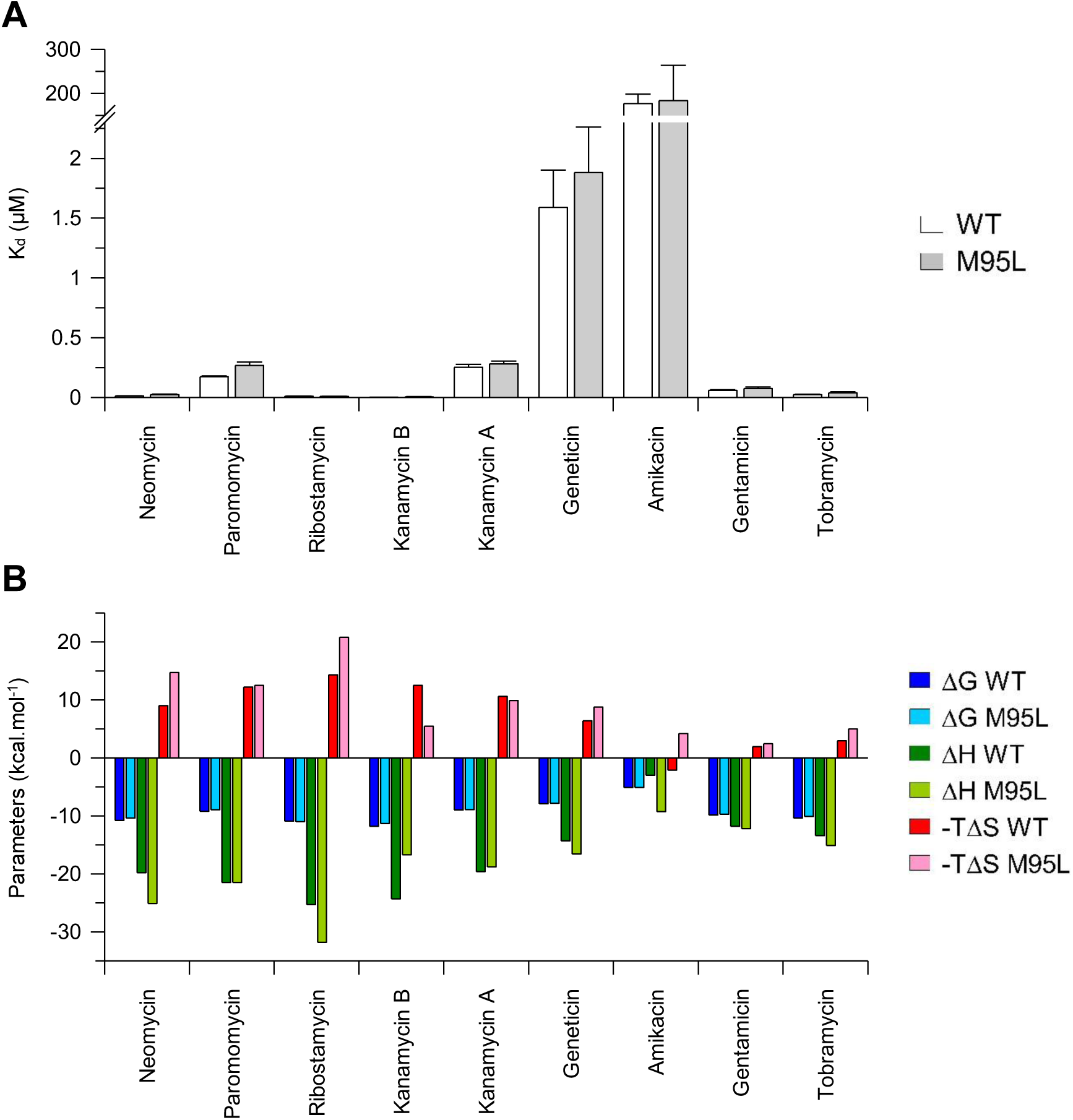
Dissociation constants and thermodynamic parameters for the binding of different aminoglycosides to WT or M95L APH(3’)-IIb. (**A**) K_d_ for aminoglycosides with APH(3’)-IIb WT (white bars) or the M95L variant (grey bars). (**B**) Thermodynamic parameters for aminoglycoside binding to APH(3’)-IIb WT (dark colors) or to the M95L variant (light colors). The calculated Gibb’s free energy (ΔG) is shown in blue, the binding enthalpy (ΔH) in green and the binding entropy (–TΔS) in red. Values of K_d_, stoichiometry (N) and thermodynamic parameters are given in **S2 Table** and binding isotherms are shown in **S2 Fig**.

Now, when comparing 4,6-disubstituted aminoglycosides, the presence of a 2’-amine in kanamycin B appears to be much more favorable than the hydroxyl in kanamycin A (yellow circle in **Table 1**). This difference alone between the two aminoglycosides seems to have an impact on the catalytic rate constant (k_cat_), but little on the affinity (K_d_). The k_cat_ values are comparable between kanamycin B and geneticin, which both have a 2’-amine. However, the K_m_ and K_d_ of geneticin are larger than those of kanamycin B, certainly due to the presence of additional methyl groups (orange boxes in **Table 1**). The same applies to amikacin, which has an L-HABA group linked to the N1, compared with kanamycin A. Finally, gentamicin and tobramycin without 3’-hydroxyls (red circles) are poor substrates for the enzyme, although they are good ligands.

### SAXS analysis predicts a monomeric form for APH(3’)-IIb

Wild-type APH(3’)-IIb was analyzed by SAXS in the absence or presence of the reducing agent, 1 mM DTT. In both cases, the curves obtained show that the protein is present in soluble, non-aggregated form up to 10 mg/mL (**S3 Fig**). Bayesian interference analysis predicts a molecular weight between 35.8 and 39.8 kDa in the absence of DTT, and between 34.2 and 37.3 kDa in the presence of DTT. As the expected molecular weight of APH(3’)-IIb with its 6-His tag is 32 kDa, we can conclude that the enzyme is predominantly monomeric in presence or not of reducing agent. The absence of aggregates allows us to envisage its crystallization.

### APH(3’)-IIb WT and M95L in apo form have a similar crystal structure

We have identified different crystallization conditions for the two APH(3’)-IIb variants. APH(3’)-IIb WT crystallized between 9 and 12 mg/mL in 0.1 M HEPES pH 7.4; 0.6 M tri-sodium citrate and 0.025 M KCl supplemented with microcrystals. APH(3’)-IIb M95L crystallized at 9 mg/mL in 20% PEG3350 and 0.3 M MgCl_2_ supplemented with microcrystals.

We determined the structure of both variants by molecular replacement using the crystal structure of *K. pneumoniae* APH(3’)-IIa (PDB 1ND4), which has 53% sequence identity with APH(3’)-IIb. The structures of APH(3’)-IIb WT and the M95L variant were solved at a resolution of 1.79 Å and 2.37 Å respectively (PDB in progress, **S4 Table**). Unsurprisingly, the structures of the two variants are identical with exception of the mutated gatekeeper residue at position 95 (**Fig 5**). The overall structure of these enzymes is, as expected, similar to that of the other members of the APH family. It may be noted that, as with other known APH structures (e.g. PDB 6FUC, 3CSV and 3DXP), the loop between the sheets β1 and β2 is poorly defined, with seven residues of this loop absent from the electron density in both the WT and M95L forms of APH(3’)-IIb. This characterizes the high flexibility of this loop, which can be found in different conformations as illustrated in **S4 Fig**.

**Fig 5.**
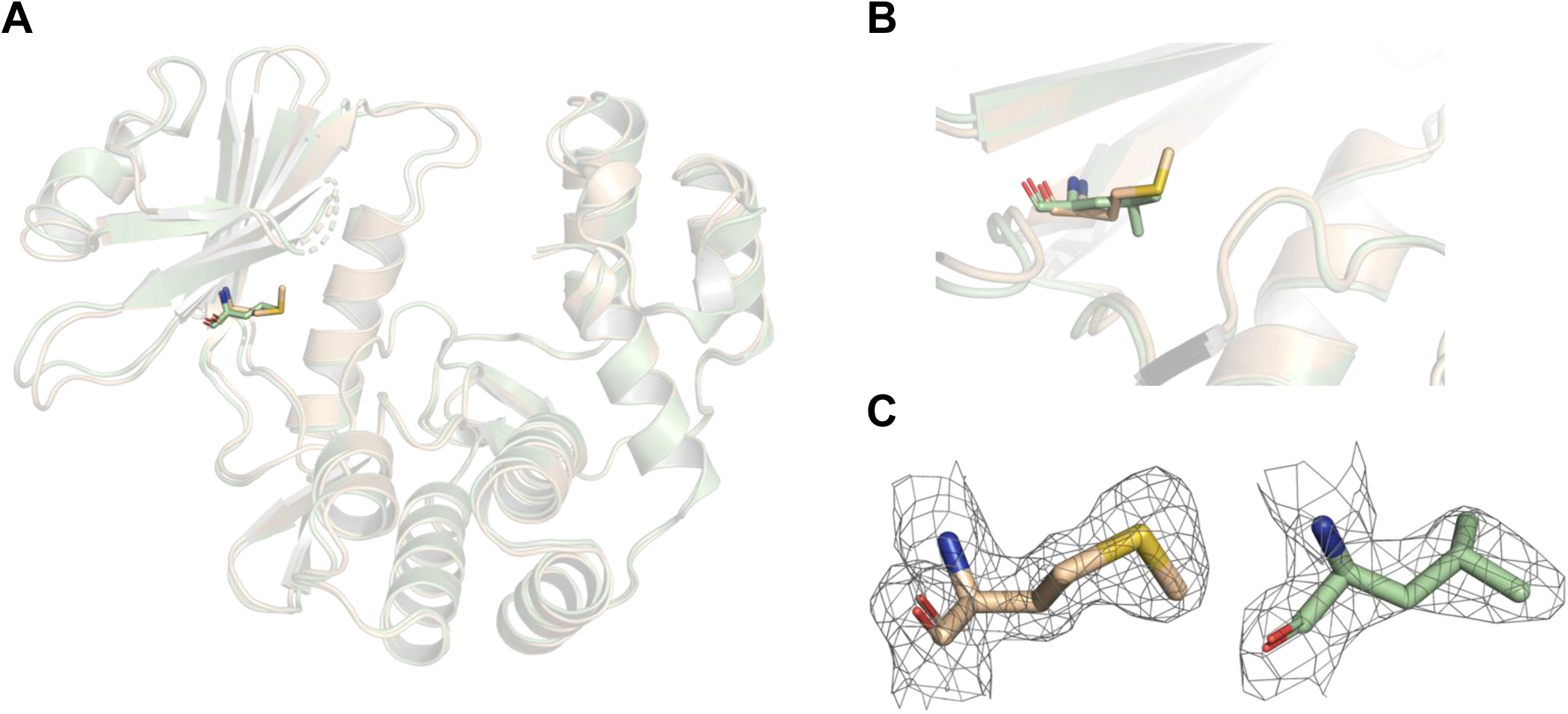
Crystal structures of the WT and M95L APH(3’)-IIb apoenzymes determined at 1.79 and 2.37 Å respectively. (**A**) Superposition of the two crystal structures. (**B**) Zoom on the superposition of the mutated gatekeeper. (**C**) Omit map of the gatekeeper residues, a methionine in the case of the wild-type enzyme, and a leucine in the case of the variant, contoured at a sigma level of ± 1 in the crystal structure. The WT and M95L structures are colored in wheat and pale green, respectively.

The electron density of the gatekeeper residues is well defined in the crystal structures, leaving no doubt as to the nature of these two amino acids (**Fig 5C**). As we have characterized differences in the activity and the affinity of these two enzymes for their natural ligands, we wondered how these results could be explained structurally. The higher electronegativity of the methionine sulfur atom may be responsible for the greater affinity of the wild-type enzyme for MgATP, compared with the M95L variant. Indeed, this sulfur atom could interact via hydrogen bonds resulting in the stabilization of the nucleobase into the catalytic pocket, whereas the leucine in the variant would not show any hydrogen bonds. However, these apoenzyme structures alone do not allow further interpretation.

### Transfer of ATP γ-phosphate to the aminoglycoside can occur in the crystal

We have determined the structure of WT APH(3’)-IIb in complex with the aminoglycoside geneticin (**S4 Table**). The electron density of this ligand is moderately well characterized, allowing us to place the three rings as well as certain hydroxyl and amine groups. The exact position of some groups is still unclear, although the interacting residues are clearly identified.

We also determined the structure of the M95L variant of APH(3’)-IIb in complex with MgADP and 3’-phosphorylated kanamycin A, PhosKanaA (**S4 Table**). It was interesting to see that the enzyme was active, even in the crystal, as we soaked MgATP and kanamycin A. To our knowledge, this is the first time that the phosphotransfer has been observed in a crystalline form of APH. This crystal structure therefore represents the first aminoglycoside kinase structure with a bound phosphorylated antibiotic substrate. The electron density of both ligands is well defined in the structure. However, the electron density around the phosphate transfer region is noisier, as we could imagine a bridge between ATP and kanamycin A. This can be explained by the fact that multiple events are compiled to obtain this electron density and that in each event the phosphate transfer state is probably different.

As in other APHs, the ADP nucleotide is located in the N-terminal domain, and the interactions with the protein are conserved (**S4 Fig**). The backbone of two residues of the hinge, S96 and V98, interact via hydrogen bonds with the N3 and N2 of adenine, respectively. Two hydrophobic residues, F49 and I212, participate in π-stacking and Van der Waals interactions with the adenine. The catalytic triad is composed of K51, E64 and D213, involved in the transfer of the γ-phosphate of ATP to aminoglycoside. Finally, two magnesium ions stabilize the phosphate groups and interact with the catalytic D213 and with two water molecules.

As expected, the cavity housing the aminoglycosides, which comprises a number of amino groups on their sugar rings, is predominantly acidic. Eight of the eleven interacting residues are acidic: D162, D164, D165, D195, D213, D232, D265 and E266. They interact with PhosKanaA through hydrogen bonds. In addition, R216 and R231 also participates to hydrogen bonding of PhosKanaA. Finally, F268 interacts via its backbone with the amine at position 6’.

## Discussion

### The nature of the gatekeeper residue explains the specificity of APH(3’)-IIb for ATP

Several studies have already identified that, in comparison to Ser/Thr protein kinases, the nature of a gatekeeper residue is decisive for the nucleotide specificity of APHs [35–39]. It is now accepted that when the gatekeeper residue is a methionine, as in APH(3’)-IIIa, ATP binding is favored by its hydrogen-bonding interaction with adenine N6. When this residue is a phenylalanine, as in APH(2’’)-IVa, the enzyme can bind either ATP or GTP. In the case of GTP, the base is displaced and stabilized by a network of highly ordered water molecules. Finally, a bulky tyrosine gatekeeper, as in APH(2’’)-IIIa, prevents ATP binding and promotes GTP binding via a hydrogen bond with guanine N7.

Here, in wild-type APH(3’)-IIb, the gatekeeper residue is a methionine, so it’s not surprising that the enzyme is ATP-specific. In a previous study, Boehr *et al.* generated mutants of the gatekeeper residue of APH(3’)-IIIa as either leucin or alanine [38]. They found that the alanine mutation had no major impact on the K_m_ or K_d_ for ATP. On the other hand, the leucine mutation, like the one we found spontaneously in our protein, had a greater impact: 3.6-fold increase in K_m_ for ATP and 4.7-fold increase in K_d_ for adenosine. This correlates well with the decreases in K_m_ for ATP (8-fold) and K_d_ for ADP (1.65-fold) or ATP (2.75-fold) that we measured with the M95L mutant compared with wild-type APH(3’)-IIb.

The spontaneous gatekeeper mutation we encountered during the cloning process could be of interest to bacteria, given that this specific amino acid is a key residue for the enzyme’s activity, and that in APH(3’)-IIa from *K. pneumoniae*, the nature of the gatekeeper is a leucine, which is the same amino acid as that resulting from the mutation in the APH(3’)-IIb variant.

### The broad active site of APH(3’)-IIb explains its low specificity for aminoglycosides

Comparison of the crystal structures of APH(3’)-IIb M95L ADP PhosKanA and APH(3’)- IIa from *K. pneumoniae* in complex with kanamycin A (PDB 1ND4, [40]) reveals that the eleven residues interacting with kanamycin A are conserved between the two enzymes, except for the D165 in APH(3’)-IIb that is replaced by E160 in APH(3’)-IIa (**Fig. 7A** and **B**). The salt bridge between R226 and D261 in APH(3’)-IIa is also conserved in APH(3’)-IIb between R231 and D265. Comparison with the aminoglycoside binding site in the structure of APH(3’)-IIIa in complex with ADP and kanamycin A (**Fig. 7C**)shows conservative substitutions from D to E (D162, D164 and D165 to E157, E160 and E161) and the substitution of D232 into S227, which means that this residue no longer interacts directly with kanamycin A, but via a water molecule. Unlike APH(3’)-IIIa, which shows a conformational change in the loop containing residues 150 to 165, termed the aminoglycoside-binding loop, that closes the site upon aminoglycoside binding [41], the APH(3’)-IIa and -IIb sites do not undergo such a large conformational change (**Fig. S4B and C**) and remain more accessible, as illustrated in **Fig. 7D-F**. This may explain the higher affinities measured by ITC with APH(3’)-IIb (**S2 Table**) than with APH(3’)-IIIa [42].

**Fig 6.**
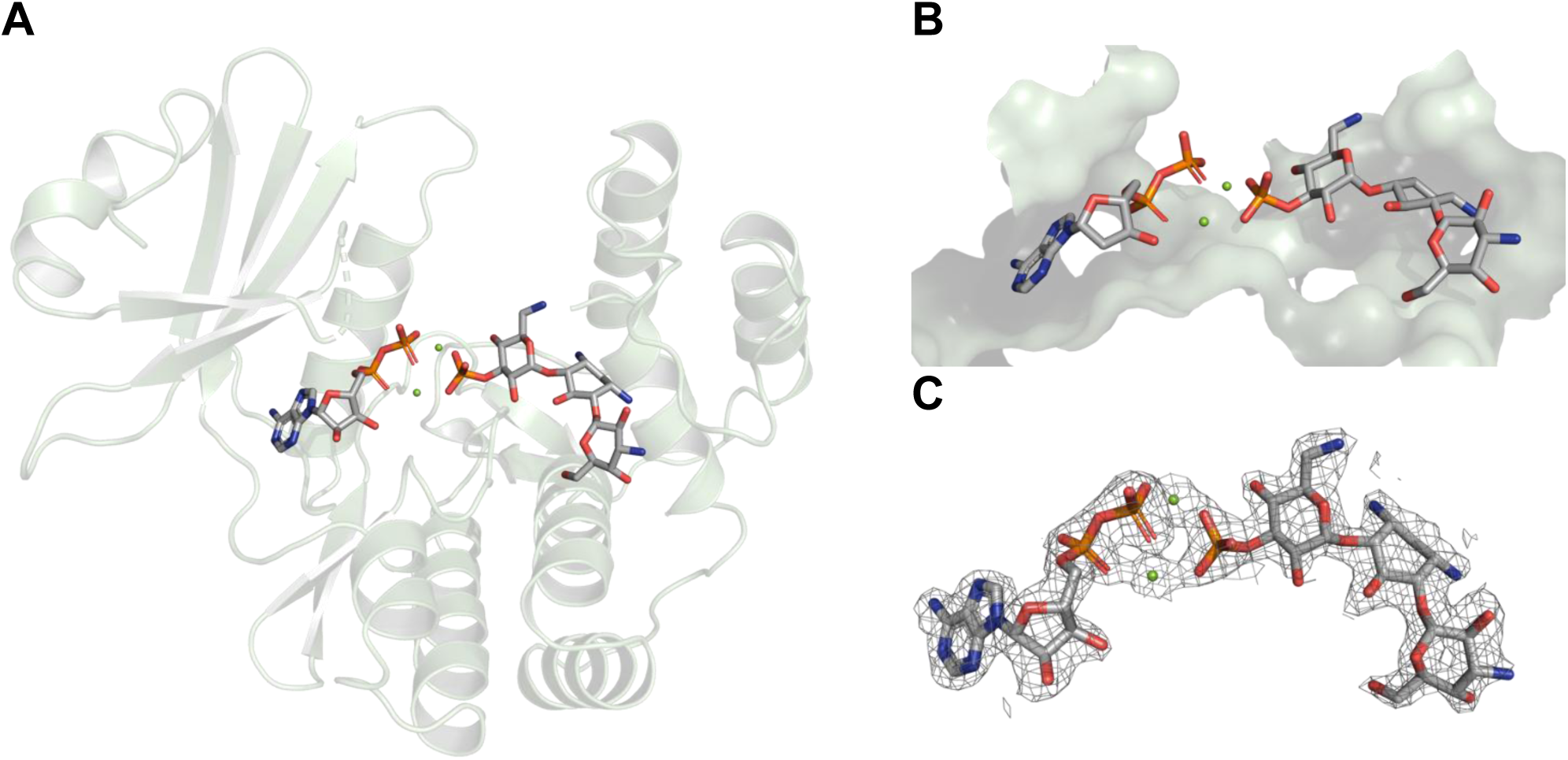
Crystal structure of the M95L APH(3’)-IIb enzyme in complex with MgADP and phosphorylated kanamycin A determined at 2.01 Å. (**A**) Overview of the ternary complex APH(3’)-IIb M95L ADP PhosKanA crystal structure. (**B**) Zoom on the binding site cavities of APH(3’)-IIb M95L where the two ligands are located. (**C**) Omit map of ADP, and PhosKanaA, contoured at a sigma level of ± 1 in the crystal structure.

**Fig 7.**
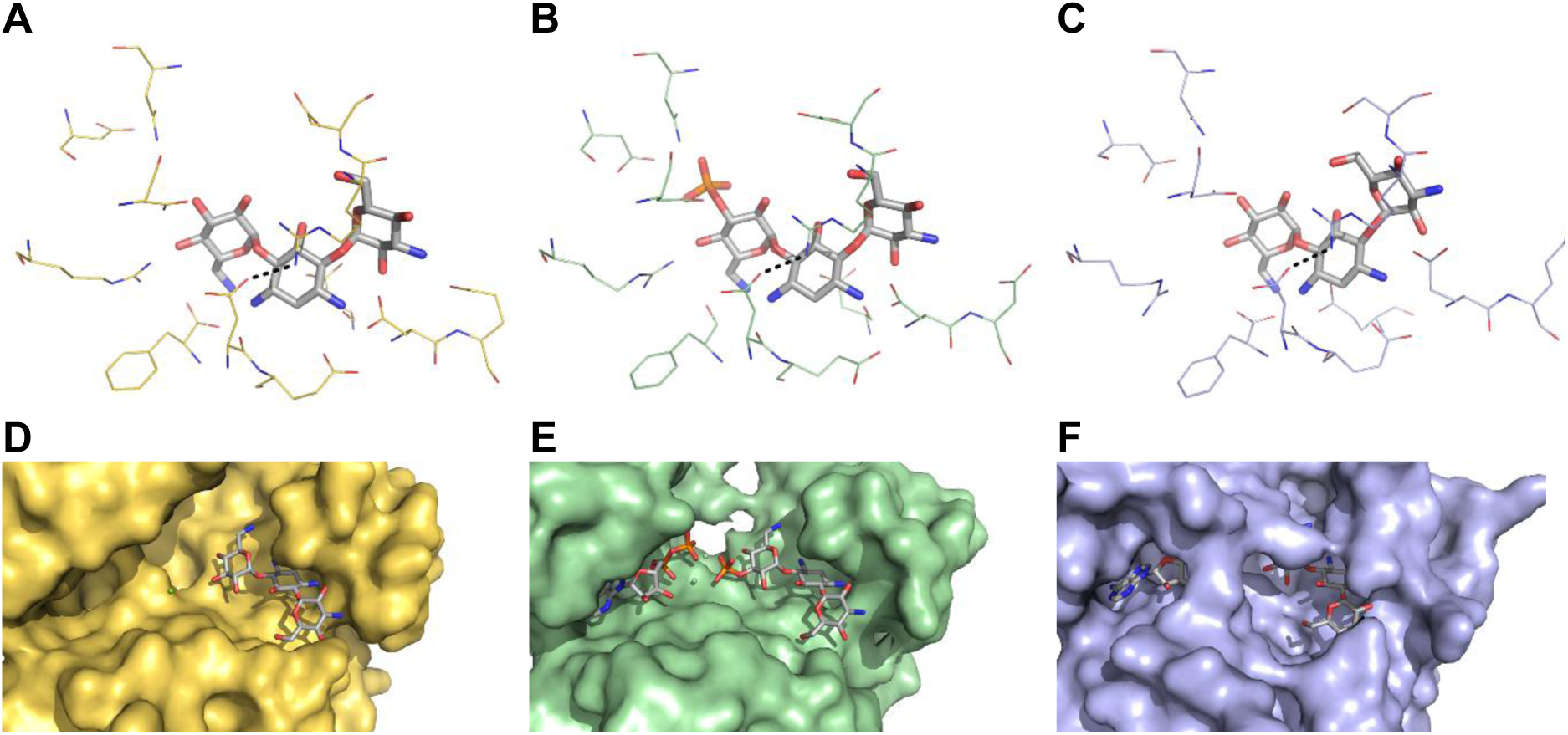
Comparison of the aminoglycoside binding sites of APH(3’)-IIa, APH(3’)- IIb and APH(3’)-IIIa. The aminoglycoside, kanamycin A 3’-phosphorylated or not, are shown as grey sticks. Residues interacting with the aminoglycoside in (**A**) APH(3’)-IIa, (**B**) APH(3’)-IIb or (**C**) APH(3’)-IIIa are shown as lines, and the salt bridges as dotted lines. Surface representations of the active site of (**D**) APH(3’)-IIa, (**E**) APH(3’)-IIb or (**F**) APH(3’)-IIIa are shown in yellow, pale green and pale blue respectively with the ligands in grey sticks.

In a previous study, we showed that with APH(2’’)-IVa, there was a correlation between the molecular weight (or number of non-hydrogen atoms) of the aminoglycosides and the K_d_ value: the larger the aminoglycosides, the lower the affinity [43]. The high accessibility of the APH(3’)-IIb aminoglycoside binding site is consistent with the lack of such a correlation. So, unlike APH(2’’)-IVa, APH(3’)-IIb can also bind 4,5-disubstituted aminoglycosides, with 3 or 4 cycles. We tried soaking APH(3’)-IIb crystals with 4,5-disubstituted aminoglycosides, but all the structures obtained were empty. Aminoglycoside docking is not easy with aminoglycosides either. So, in the absence of structural data, we placed these ligands in the electron density corresponding to PhosKanA. This manual positioning of the 4,5-disubstituted aminoglycosides shows that they could fit perfectly into the enzyme’s active site and create potential additional interactions (highlighted by green circles in the **S5F-G Fig.**), in line with their high affinity. This will have to be validated by obtaining the structure of the complexes, for example by co-crystallization.

Now, if we compare kanamycins A and B, which differ only by the nature of the group in the 2’ position (yellow circles in **Table 1** and in **S5B-C Fig.**), we notice that the amine of kanamycin B is more favorable than the hydroxyl of kanamycin A for both affinity (**S2 Table**) and catalysis (k_cat_, **Table 1**). As mentioned previously [42], the pKa of the 2’-amino group of kanamycin B is ≈8.6 and so, at pH 7.5, the 2’-amino group would be ≈90% protonated, while all the others would be ≈50% or less protonated. We suggest that the presence of a 2’-amine would not initiate additional interactions with the enzyme compared to a 2’-hydroxyl, but that the amine’s positive charge could be favorable to phosphate transfer to the 3’-hydroxyl. With the exception of gentamicin and tobramycin for reasons discussed below, with all the aminoglycosides with an amine at this position tested (neomycin, paromomycin, ribostamycin, kanamycin B and geneticin), the k_cat_ value measured was similar and higher (between 1.98 and 2.64 s^-1^) than that of kanamycin A (0.58 s^-1^). This observation could be useful for the development of new aminoglycosides that would be less favorably inactivated by this enzyme.

As for tobramycin and gentamicin, which have no 3’-hydroxyl (red circles in **Table 1** and in **S5H-I Fig.**), they are, as expected, very poor substrates despite high affinities (K_d_ of 0.023 and 0.059 µM, respectively). However, while no activity could be measured with tobramycin, a weak activity seems to exist with gentamicin. Using the same manual placement in the electron density of 3’-phosphokanamycin A as before, we suggest that gentamicin may bind backwards and potentially be phosphorylated on the hydroxyl in the 4’’ position (black circles in **Table 1** and in **S5I Fig.**). As its position is not optimal in relation to ATP γ-phosphate, this could explain the low activity mesured.

Further experiments are needed to explain how amikacin binding occurs in the binding site of wild-type and M95L variants of APH(3’)-IIb, but also on other members of the APH family. Determining the structure of these enzymes in complex with amikacin could help us understand the binding mechanism of this antibiotic. However, several attempts to resolve this complex were unsuccessful, as no density was visible for amikacin in the aminoglycoside binding site of the protein, irrespective of soaking time (from 10 min to 1 day). In addition, a noisier density was observed in the C-terminal part of APH(3’)-IIb from crystals soaked with this antibiotic, indicating that amikacin binding could disrupt the integrity of the aminoglycoside binding site of the protein. In the meantime, we have tried to place amikacin in the electron density of PhosKanA (**S5J Fig.**). When the three rings of amikacin are placed at the same positions as those of kanamycin A, the L-HABA group at the N1 position of amikacin clashes with residues D162, F163 and D164 (purple square in the figure).

### Considerations for the design of specific APH inhibitors

Time-resolved crystallography could be of great help in visualizing the transfer of γ-phosphate from ATP to aminoglycoside molecules. Indeed, this method could help us to understand amino acid dynamics at the catalytic site and allosteric changes occurring during the reaction, and identify reaction intermediates that could be targeted [44]. Combined with a classical fragment-based approach to drug design, it could help design competitive or allosteric inhibitors of this enzyme. Similar to what has been done with β-lactamase inhibitors, the development of inhibitors of APH(3’)-IIb, and to a greater extent of the APH family, could help counter aminoglycoside resistance and re-sensitize bacteria resistant to this class of antibiotics. This opens up a hopeful avenue in the fight against antibiotic resistance, given that aminoglycoside antibiotics are widely used as the last therapeutic barrier against multi-resistant bacterial infections.

## Materials and Methods

### Cloning the *aph(3’)-IIb* gene and transforming bacteria

The *aph(3’)-IIb* gene was identified using the UniProtKB database (Q9HWR2_PSEAE). The corresponding sequence was extended by 5’- CGCGCGGCAGCCAT and GGATCCGGCTGCTAAC-3’ and optimized for expression in *E. coli* (Integrated DNA Technologies GMBH, Germany) for subsequent cloning into plasmid pET-15b using the In-Fusion^®^ HD Cloning Kit (Takara Bio USA, Inc.). Prior to the reaction, purified pET-15b was incubated for 1 h at 37 °C with the restriction enzymes NdeI and BamHI for digestion. The digestion product was loaded onto a 1% agarose gel, migrated for 30 min at 100 V, and the linearized pET-15b was then purified from the agarose gel using the PCR clean-up kit (Macherey-Nagel, France). The synthetic gene (IDT technologies) was resuspended in 40 µL of sterile water. The In-Fusion reaction was then performed by mixing linearized pET-15b with *aph(3’)-IIb* gene and incubating this reaction mix in a thermocycler at 50 °C for 15 min.

The digestion product was loaded onto a 1% agarose gel, migrated for 30 minutes at 100 V, and the linearized pET-15b was then purified from the agarose gel using the Macherey-Nagel PCR clean-up kit. The genes were resuspended in 40 µl of sterile water. The In-Fusion reaction was then performed by mixing linearized pET-15b with the *aph(3’)-IIb* gene and incubating the reaction mixture in a thermocycler at 50°C for 15 min.

The In-Fusion cloning product was used to transform both *E. coli* DH5α and BL21 DE3. Aliquots of chemo-competent cells were gently thawed on ice. A 2 µL volume of the cloning product was incubated with 50 µL of bacteria. The bacteria were then heat-shocked at 42 °C for 45 s and then directly kept on ice for 2 min. The transformed bacteria were supplemented with 300 µL of Luria-Bertani (LB) medium and incubated at 37 °C for 1h under orbital shaking. 200 µL of bacteria were spread on an LB agar plate supplemented with ampicillin. The plates were then incubated overnight at 37 °C. The following day, three to six clones per plate were picked and plated onto a new LB agar plate, supplemented with ampicillin or ampicillin and chloramphenicol, or used to inoculate 5 mL of liquid LB with the corresponding antibiotics for selection. Plates and liquid cultures were incubated overnight at 37 °C with orbital shaking for liquid cultures. Plasmids were extracted from 5 mL of DH5α cultures using the Macherey-Nagel Plasmid kit protocol except for the elution volume which was reduced from 50 µL of Elution Buffer (protocol) to 30 µL of sterile water. The restriction profiles of the extracted plasmids were then checked before being sent for sequencing to Eurofins, France.

A plasmid containing the correct *aph(3’)-IIb* gene was used to transform BL21 DE3 bacteria as described above for DH5α.

### Expression and purification of recombinant APH(3’)-IIb

Recombinant APH(3’)-IIb was expressed using the pET15b plasmid described above in *E. coli* BL21 DE3. Bacterial glycerol stocks made after sequencing were used to purify the enzyme. First, the glycerol stock was scraped and plated on an ampicillin-supplemented LB agar plate to select the clones possessing the plasmid encoding the protein of interest, then incubated overnight in a static incubator at 37 °C. On the second day, a clone was peaked and used to inoculate a 500 mL flask containing 200 mL of LB supplemented with ampicillin and chloramphenicol. This pre-culture was incubated overnight at 37 °C with orbital shaking.

On the third day, the pre-culture was used to produce 3 L of culture in four flasks of 2 L each of them containing 750 mL of LB supplemented with ampicillin. To do this, the optical density at 600 nm (OD_600_) of pre-culture was measured and the volume of pre-culture required to reach an initial OD_600_ of 0.1 in each flask. The culture was then grown at 37 °C with an orbital shaking to reach an OD_600_ between 0.6 and 0.8 before induction. Once the culture had reached this OD_600_, induction of protein expression was initiated by adding 1 mM of IsoPropyl-Thio-β-Galactoside (IPTG) to each flask. After addition of IPTG, the flasks were directly moved from 37 °C to 18 °C and incubated overnight with an orbital shaking. The overnight culture was then centrifuged for 30 min at 6,000 rpm and 8 °C to harvest the bacteria. The pellet was then resuspended in 100 mL of Buffer A supplemented with two tablets of cOmplete™ protease inhibitor cocktail, 10 µg/mL of Lysozyme and 1 µg/mL of DNase. The resuspended pellets were pooled and sonicated on ice with a probe for 10 min at an intensity of 50 and 2-second pulses.

The lysate was then centrifuged during 30 min at 18,000 rpm and 8 °C to separate soluble fraction from cellular remains and insoluble components. The supernatant was then filtered on 5 µm followed by 0.45 µm filters prior to the affinity chromatography. Two 5 mL HisTrap FF Crude columns were connected in tandem on an Äkta Pure equilibrated with 10 column volume (CV) of Buffer A (50 mM NaH_2_PO4, 300 mM NaCl, 10 mM Imidazole, 1 mM DTT, pH 8). The filtered supernatant was injected on columns using a sample pump equipped with an air sensor to avoid formation of air bubbles. Once all the supernatant was injected, the columns where washed with 15 CV of Buffer A. Elution of the protein was performed using a linear gradient of Buffer B (50 mM NaH_2_PO4, 300 mM NaCl, 500 mM Imidazole, 1 mM DTT, pH 8) going from 0% to 5% for 2 CV in a first time. Then, a step was performed at 5% and maintained for 3 CV. These two first steps allowed to exclude a contaminant from the protein of interest. The elution gradient was then pursued with a linear gradient from 5% to 50% for 5 CV and the protein was eluted at approximately 12% of Buffer B and collected in fractions of 2 mL. After reaching the 50%, the gradient was set to a step at 100% of Buffer B to elute every possible particle that could have been stuck on the columns. The columns where then washed with 10 CV of water, then 10 CV of ethanol 20% and stored at 4 °C.

The fractions corresponding to the peak of elution of the protein of interest were analyzed on Tris-glycine 4-12% acrylamide precast gel from Invitrogen™. Fractions of interests were pooled and divided in two to performed two separate size exclusion chromatography respectively in Tris (50 mM Tris, 40 mM KCl, 1 mM MgCl_2_, 1 mM DTT, pH 7.5) or HEPES (50 mM HEPES, 40 mM KCl, 1 mM MgCl_2_, 1 mM DTT, pH 7.5) buffer. These buffers were chosen for optimal protein stability previously determined by Thermal Shift Assay [45].The two pools were concentrated in 15 mL Amicon® Ultra 30 kDa cut-off concentrator until an injectable concentration as well as volume of protein for size exclusion chromatography. The concentrated protein was loaded on either a HiLoad™ 16/60 or 26/60 Superdex™ 75 prep grade from GE Healthcare. After the elution, the fractions of interest were analyzed on Tris-glycine 4-12% acrylamide precast gel from Invitrogen™. The fractions of interest were pooled and concentrated to a concentration > 15 mg/mL Tris buffer and > 500 µM in HEPES buffer. Proteins were aliquoted and stored at –80 °C.

### Enzymatic activity measurements

Measurements of the enzymatic activity of APH(3’)-IIb were performed either using an enzyme coupled system or by stopping the reaction at different times and quantifying reaction products by HPLC. Using the enzyme coupled system, the experiments were carried out in triplicate at 25 °C in the Tris buffer using the pyruvate kinase and lactate dehydrogenase coupled system, as previously described [23]. The final reaction mixture contained 25 nM APH(3’)-IIb, 140 µM NADH, 2 mM PEP, 1× LDH/PK and either 100 µM Kanamycin A and various concentrations of MgATP or MgGTP, or 100 µM MgATP and various concentrations of aminoglycosides. All chemicals and enzymes were from Sigma-Aldrich.

### Assay of NADH, nucleotide and aminoglycoside concentrations

As commercial NADH, nucleotide and aminoglycoside powders contain a sometimes unknown quantity of ions and water molecules, the effective concentrations of prepared stock solutions were measured. NADH and nucleotide concentrations were measured using a NanoPhotometer^®^ NP80 (Implen) using a molar extinction coefficient of 6.22 at 340 nm for NADH, 15.4 at 259 nm for adenosine-based nucleotides and 13.7 at 253 nm for guanosine-based nucleotides.

Aminoglycoside concentrations were determined using the coupled enzymatic system described above. In this case, the final concentrations were 1 µM APH, 350 µM MgATP, 200 µM NADH, 2 mM PEP, 1x LDH/PK and 0 to 140 µM (theoretical) aminoglycosides. Measurement of the concentration of NADH consumed at the end of the reaction, when all the aminoglycosides have been phosphorylated, enables us to plot a standard range for determining the actual concentrations of aminoglycosides. In the case of aminoglycosides not substrates of APH(3’)-IIb, recombinant APH(2’’)-IVa was used (purified as in [43]).

### Affinity measurements

Experiments were performed at 25 °C in HEPES buffer using a MicroCal iTC200 or PEAQ-ITC device (Malvern Panalytical). The enzymes were used at a final concentration of 120 μM in the ITC cell and the ligands at 2-5 mM in the syringe. Titrations consisted of 19 injections of 2 μL of ligand solution or 25 injections of 1.5 μL of ligand solution every 150 s with a stir speed of 750 rpm. K_d_ values were determined using the MicroCal PEAQ-ITC analysis software version 1.41.

### SAXS

Data were collected on beamline BM29 at ESRF (doi.org/10.15151/ESRF-ES-1215821670). APH(3’)-IIb was used in batch at a concentration of 2, 4 or 8 mg/mL in Tris buffer with or without 1 mM DTT. Data were scaled, averaged and analyzed using primus/qt ATSAS 3.0.3 software.

### Crystallogenesis and structure determination

APH(3’)-IIb M95L crystallized between 9 mg/ml in 0.3 M MgCl2 and 20% PEG 3350 coupled to micro seeding with microcrystals of the protein generated during the optimisation of the conditions. APH(3’)-IIb wild-type crystallized between 9 and 12 mg/mL in 0.1 M HEPES pH 7.4, 0.6 M tri-Sodium Citrate and 0.025 M KCl coupled to micro seeding with microcrystals of the protein generated during the optimisation of the conditions.

Both proteins were crystallized using the hanging drop vapor diffusion method, mixing 1 µl of purified protein with 1 µl of reservoir solution. Diffraction data were collected for APH(3’)-IIb WT and M95L respectively to 1.79 Å and 2.37 Å resolution at the European Synchrotron Radiation Facility (ESRF). The X-ray structures of the apoenzymes were determined by molecular replacement using the structure of APH(3’)-IIa from *K. pneumoniae* (PDB 1ND4) solved in complex with kanamycin A as the search model.

The structures of the enzyme in complex with ligands were obtained by soaking apoenzyme crystals with the corresponding ligands. The ternary complex APH(3’)-IIb M95L ADP PhosKanA structure has been generated by soaking 2.4 mM of MgATP for 22 min and in the same drop, 0.5 mM of kanamycin A was added 11 min after MgATP and left for 10 min before the crystal was frozen in liquid nitrogen. The complex APH(3’)-IIb WT Geneticin structure has been generated by soaking 0.4 mM of geneticin for 9 min before the crystal was frozen in liquid nitrogen.

Ligand restraints were generated with the Grade Web Server 2.0.9. The atomic models were rebuilt in Coot 0.9.8.94 software from the CCP4-9 Software Suite and refined using Phenix.Refine from Phenix 1.21.1 software. Data collection and refinement statistics for the four structures are summarized in **S4 Table**. Figures were generated with PyMOL 1.8.4.2 (http://pymol.org/).

## Acknowledgments

This work was supported by ANR SIAM (ANR-19-AMRB-0001), the Association Vaincre la Mucoviscidose (RF20220503015), and the Association Grégory Lemarchal. The CBS is part of the ChemBioFrance research infrastructure and is a member of the FranceBioImaging (FBI) and the French Infrastructure for Integrated Structural Biology (FRISBI), two national infrastructures supported by the Agence Nationale de la Recherche (ANR-10-INBS-0004 and ANR-10-INBS-0005, respectively). We would like to thank Dr. Assia Mouhand and Dr. Pau Bernado of CBS for their help in analyzing of SAXS data and the personal of ESRF for the acquisition of SAXS and X-ray diffraction data.

