## Supplementary Tables and Figures for "Structural and molecular basis of resistance to aminoglycoside antibiotics by APH(3’)-IIb from *Pseudomonas aeruginosa*"

**Short title:** Characterization of *Pseudomonas aeruginosa* APH(3')-IIb

**Julien Kowalewski[1], Mathilde Tomaszczyk[1], Jean-François Guichou[1], Muriel Gelin[1], Gilles Labesse[1], Corinne Lionne[1]***

[1] Centre de Biologie Structurale (CBS), CNRS UMR 5048, University of Montpellier, INSERM U 1054, Montpellier, France.

*

### Supplementary files

**S1 Table. Dissociation constants and thermodynamic parameters for the binding of nucleotides to APH(3')-IIb.**
**S2 Table. Dissociation constants and thermodynamic parameters for the binding of different aminoglycosides to APH(3')-IIb.**
**S3 Table. General gentamicin formula and characteristics of the components in the batch used.**
**S4 Table. Data collection and refinement statistics for the four APH(3')-IIb crystallographic structures solved during this work.**
**S1 Fig. ITC isotherms with nucleotides or analogues**.
**S2 Fig. ITC isotherms with aminoglycosides**.
**S3 Fig. SAXS data**.
**S4 Fig. Comparison of crystal structures of different APHs**.
**S5 Fig. Prediction of interactions of APH(3')-IIb with 4,5- and 4,6-disubstituted aminoglycosides**.

**S1 Table. Dissociation constants and thermodynamic parameters for the binding of nucleotides to APH(3')-IIb.** $K_d$ values were calculated from ITC-derived $K_a$. The value of –TΔS were determined at 25°C. Binding isotherms and fits are given in **S1 Fig**.

| **Nucleotide** | **Wild-type** | | | | | **M95L** | | | | | |
|---|---|---|---|---|---|---|---|---|---|---|---|
| | **N** | **$K_d$** (μM) | **ΔH** ($k_{cal}$ mol$^{-1}$) | **ΔG** ($k_{cal}$ mol$^{-1}$) | **–TΔS** ($k_{cal}$ mol$^{-1}$) | **N** | **$K_d$** (μM) | **Relative to WT** (×) | **ΔH** ($k_{cal}$ mol$^{-1}$) | **ΔG** ($k_{cal}$ mol$^{-1}$) | **–TΔS** ($k_{cal}$ mol$^{-1}$) |
| MgADP | 1.02 ± 0.01 | 4.1 ± 0.5 | –9.31 ± 0.15 | –7.36 | 1.95 | 0.81 ± 0.01 | 6.7 ± 0.9 | 1.65 | –5.16 ± 0.14 | –7.07 | –1.91 |
| MgATP | 0.97 ± 0.01 | 10.0 ± 0.7 | –12.30 ± 0.20 | –6.82 | 5.44 | 1.12 ± 0.03 | 27.5 ± 4.5 | 2.75 | –3.12 ± 0.17 | –6.22 | –3.10 |
| MgAMPPCP | 0.83 ± 0.01 | 23.3 ± 1.7 | –13.80 ± 0.37 | –6.32 | 7.46 | 1 (fixed) | 66.5 ± 12.1 | 2.85 | –5.66 ± 0.50 | –5.70 | –0.04 |
| MgGDP | 1 (fixed) | 285 ± 47 | –3.01 ± 0.37 | –4.84 | –1.83 | 1 (fixed) | 238 ± 12 | 0.83 | –4.86 ± 0.17 | –4.95 | –0.09 |
| MgGTP | 1 (fixed) | 224 ± 25 | –0.60 ± 0.04 | –4.98 | –4.38 | 1 (fixed) | 245 ± 48 | 1.09 | –2.06 ± 0.28 | –4.93 | –2.87 |
| MgGMPPCP | 1 (fixed) | 196 ± 62 | –1.10 ± 0.22 | –5.06 | –3.96 | 1 (fixed) | 265 ± 33 | 1.35 | –3.61 ± 0.32 | –4.88 | –1.27 |

**S2 Table. Dissociation constants and thermodynamic parameters for the binding of different aminoglycosides to APH(3')-IIb.** MW, $N_{CNO}$, and LE represent the molecular weight, the number of non-hydrogen atoms and the ligand efficiency of aminoglycosides, respectively. $K_d$ values were calculated from ITC-derived $K_a$. The value of –TΔS were determined at 25°C. Binding isotherms and fits are given in **S2 Fig**.

| | | | Wild-type | | | | | M95L | | | | |
|---|---|---|---|---|---|---|---|---|---|---|---|---|
| **Ligands** | **MW** (g mol$^{-1}$) | **$N_{CNO}$** | **N** | **$K_d$** (µM) | **ΔH** ($k_{cal}$ mol$^{-1}$) | **ΔG** ($k_{cal}$ mol$^{-1}$) | **-TΔS** ($k_{cal}$ mol$^{-1}$) | **N** | **Kd** (µM) | **ΔH** ($k_{cal}$ mol$^{-1}$) | **ΔG** ($k_{cal}$ mol$^{-1}$) | **-TΔS** ($k_{cal}$ mol$^{-1}$) |
| Neomycin | 614.6 | 42 | 1.19 | 0.012 ± 0.002 | -19.8 ± 0.3 | -10.8 | 9.0 | 0.81 | 0.023 ± 0.006 | -25.1 ± 0.4 | -10.4 | 14.7 |
| Paromomycin | 615.6 | 42 | 0.91 | 0.173 ± 0.008 | -21.5 ± 0.1 | -9.2 | 12.2 | 0.73 | 0.267 ± 0.029 | -21.5 ± 0.4 | -9.0 | 12.5 |
| Ribostamycin | 454.5 | 31 | 0.80 | 0.010 ± 0.001 | -25.3 ± 0.1 | -10.9 | 14.3 | 0.61 | 0.008 ± 0.002 | -31.8 ± 0.2 | -11.0 | 20.8 |
| Kanamycin B | 483.5 | 33 | 0.95 | 0.002 ± 0.001 | -24.3 ± 0.1 | -11.8 | 12.5 | 0.98 | 0.006 ± 0.002 | -16.7 ± 0.2 | -11.3 | 5.4 |
| Kanamycin A | 484.5 | 33 | 0.89 | 0.252 ± 0.025 | -19.6 ± 0.3 | -9.0 | 10.6 | 0.89 | 0.281 ± 0.022 | -18.8 ± 0.2 | -8.9 | 9.9 |
| Geneticin | 496.6 | 34 | 1.13 | 1.59 ± 0.31 | -14.3 ± 1.0 | -7.9 | 6.4 | 0.93 | 1.88 ± 0.382 | -16.6 ± 0.9 | -7.8 | 8.8 |
| Amikacin | 585.6 | 40 | 1.00 | 177 ± 22 | -3.0 ± 0.3 | -5.1 | -2.1 | 0.82 | 184 ± 80 | -9.3 ± 4.8 | -5.1 | 4.2 |
| Gentamicin* | 477.6 | 33 | 0.93 | 0.059 ± 0.005 | -11.8 ± 0.1 | -9.9 | 1.9 | 0.93 | 0.074 ± 0.014 | -12.2 ± 0.2 | -9.7 | 2.4 |
| Tobramycin | 467.5 | 32 | 1.00 | 0.023 ± 0.003 | -13.4 ± 0.1 | -10.4 | 3.0 | 1.14 | 0.039 ± 0.008 | -15.1 ± 0.2 | -10.1 | 5.0 |

*MW and $N_{CNO}$ are given for Gentamicin C1. Characteristics of the 5 major components of Gentamicin are given in **S3 Table**.

**S3 Table. General gentamicin formula and characteristics of the components in the batch used.**

**General formula**

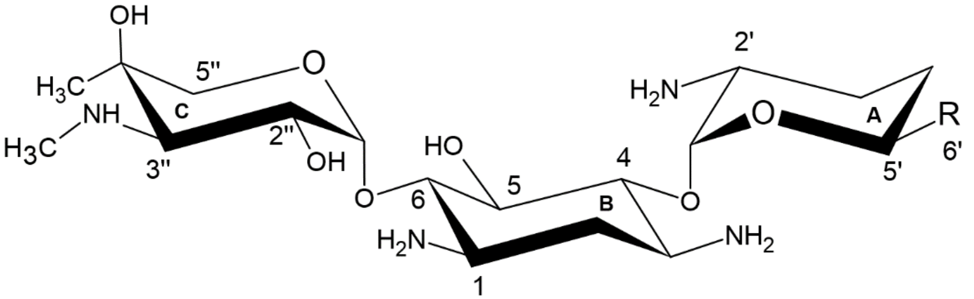


| Component | R | MW (g mol$^{-1}$) | $N_{CNO}$ | Ratio |
|---|---|---|---|---|
| C1 | 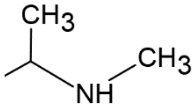 | 477.6 | 33 | C1 + C2b 33% |
| C2b | 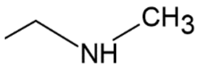 | 463.6 | 32 | |
| C1a | 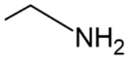 | 449.5 | 31 | C1A 20% |
| C2 | 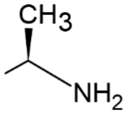 | 463.6 | 32 | C2 + C2a 47% |
| C2a | 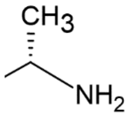 | 463.6 | 32 | |

**S4 Table. Data collection and refinement statistics for the four APH(3')-IIb crystallographic structures solved during this work.**

| | WT (PDB in progress) | M95L (PDB in progress) | M95L·ADP·PhosKanA (PDB in progress) | WT·Geneticin (PDB in progress) |
|---|---|---|---|---|
| Wavelength | | | | |
| Resolution range | 39.35 - 1.79 (1.82 - 1.79) | 90.74 - 2.37 (2.42 - 2.37) | 55.11 - 2.01 (2.04 - 2.01) | 45.01 - 2.18 (2.22 - 2.18) |
| Space group | C 2 2 21 | P 21 21 21 | P 21 21 21 | C 2 2 21 |
| Unit cell | | | | |
| a,b,c (Å) | 83.006 110.127 123.673 | 44.854 70.648 181.478 | 44.864 68.069 187.766 | 82.317 109.884 123.26 |
| α,β,γ(°) | 90 90 90 | 90 90 90 | 90 90 90 | 90 90 90 |
| Total reflections | 104670 (5059) | 48590 (3141) | 68114 (3372) | 73786 (3063) |
| Unique reflections | 100765 (4934) | 45289 (3004) | 64425 (3237) | 70683 (2976) |
| Multiplicity | 1.0 (1.0) | 1.1 (1.0) | 1.1 (1.0) | 1.0 (1.0) |
| Completeness (%) | 99.00 (93.15) | 99.56 (99.87) | 86.67 (94.02) | 99.30 (97.85) |
| Mean I/sigma(I) | 7.46 (0.98) | 6.19 (1.59) | 6.62 (1.68) | 6.34 (0.63) |
| Wilson B-factor | 22.23 | 29.81 | 26.42 | 35 |
| R-merge | 1.131e-17 (2.228e-17) | 1.11e-17 (1.467e-17) | 1.406e-17 (1.661e-17) | 1.237e-17 (3.047e-17) |
| R-meas | 1.6e-17 (3.151e-17) | 1.57e-17 (2.074e-17) | 1.989e-17 (2.349e-17) | 1.749e-17 (4.309e-17) |
| R-pim | 1.131e-17 (2.228e-17) | 1.11e-17 (1.467e-17) | 1.406e-17 (1.661e-17) | 1.237e-17 (3.047e-17) |
| CC1/2 | 1 (1) | 1 (1) | 1 (1) | 1 (1) |
| CC* | 1 (1) | 1 (1) | 1 (1) | 1 (1) |
| Reflections used in refinement | 52913 (2582) | 24373 (1572) | 36562 (1779) | 29457 (1437) |
| Reflections used for R-free | 2610 (131) | 1152 (73) | 1778 (107) | 1495 (65) |
| R-work | 0.1822 (0.3434) | 0.2013 (0.2953) | 0.1798 (0.2613) | 0.1832 (0.2914) |
| R-free | 0.2331 (0.3601) | 0.2642 (0.3591) | 0.2405 (0.2823) | 0.2373 (0.3284) |
| CC(work) | | | | |
| CC(free) | | | | |
| Number of non-hydrogen atoms | 4775 | 4241 | 4656 | 4431 |
| macromolecules | 4180 | 4047 | 4068 | 4113 |
| ligands | 1 | 24 | 154 | 49 |
| solvent | 594 | 170 | 434 | 269 |
| Protein residues | 519 | 512 | 513 | 521 |
| Nucleic acid bases | | | | |
| RMS(bonds) | 0.006 | 0.092 | 0.09 | 0.126 |
| RMS(angles) | 0.78 | 2.06 | 2.14 | 0.85 |
| Ramachandran favored (%) | 98.63 | 97.22 | 99.01 | 98.64 |
| Ramachandran allowed (%) | 1.37 | 2.78 | 0.99 | 1.36 |
| Ramachandran outliers (%) | 0 | 0 | 0 | 0 |
| Rotamer outliers (%) | 1.2 | 1.98 | 0.25 | 2.21 |
| Clashscore | 2.41 | 4.41 | 6.1 | 4.5 |
| Average B-factor | 27.29 | 34.94 | 29.4 | 38.81 |
| macromolecules | 26.53 | 35.06 | 29 | 38.62 |
| ligands | 35.99 | 44.32 | 30.68 | 55.41 |
| solvent | 32.65 | 30.71 | 32.69 | 38.65 |
| Number of TLS groups | | | | |

**S1 Fig. ITC isotherms with nucleotides or analogues**. Thermodynamics of nucleotide binding to APH(3')-IIb (**A**) WT or (**B**) M95L variants. The top panels show the differential heat released following baseline subtraction and the bottom panels show ITC binding curves.

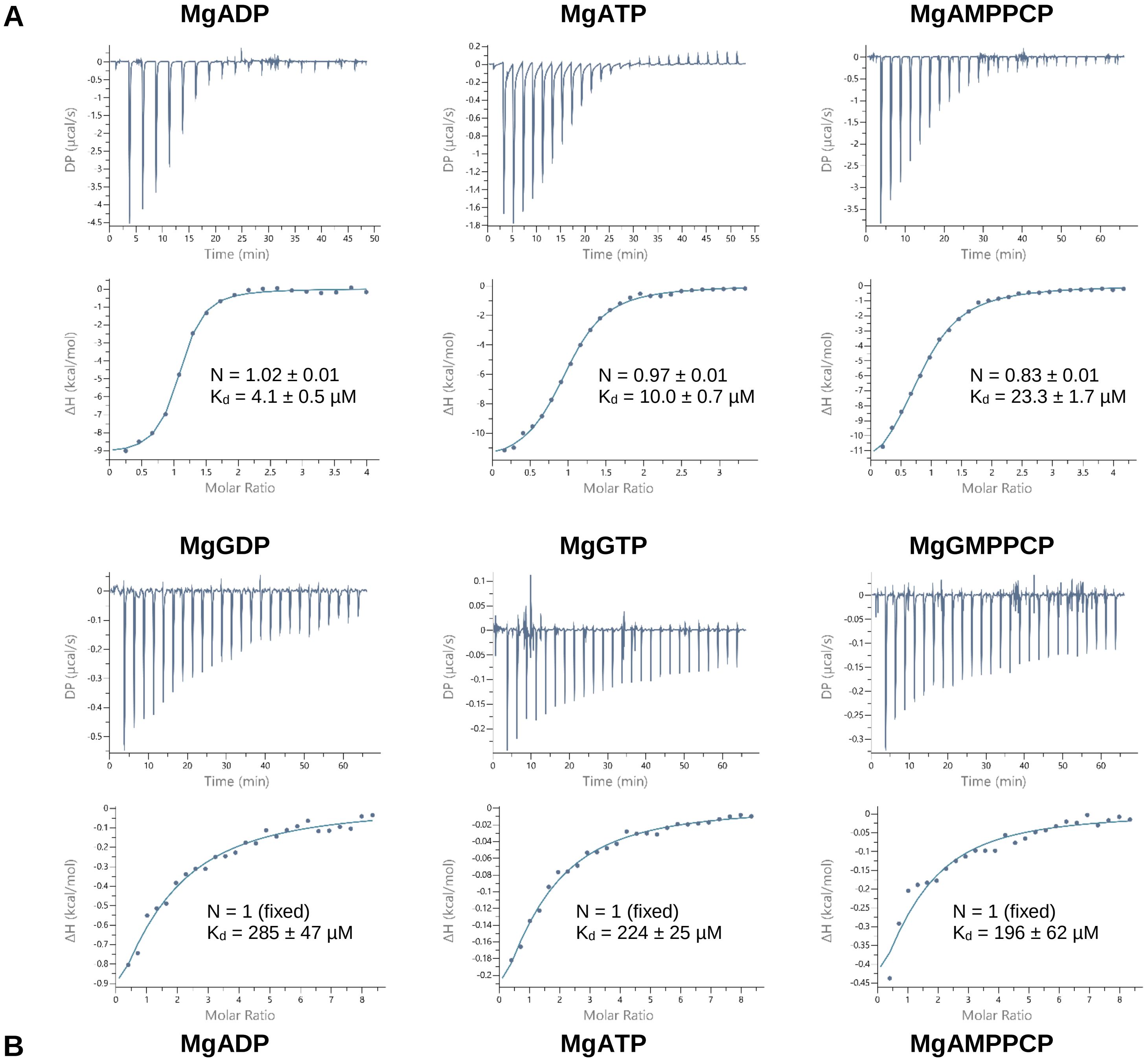

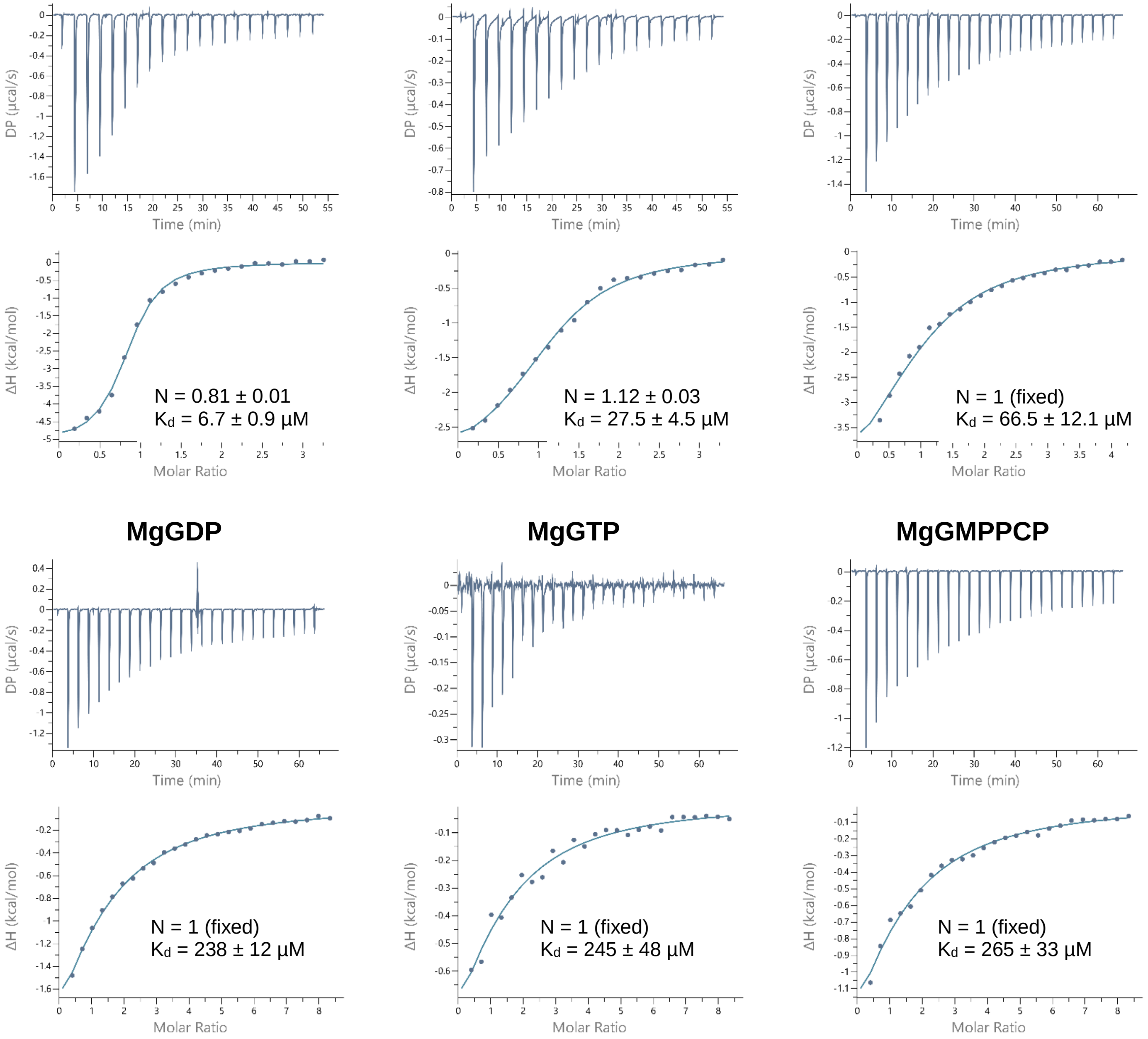
DP (µcal/s)
Time (min)
ΔH (kcal/mol)
Molar Ratio
N = 0.81 ± 0.01
Kd = 6.7 ± 0.9 µM
N = 1.12 ± 0.03
Kd = 27.5 ± 4.5 µM
N = 1 (fixed)
Kd = 66.5 ± 12.1 µM
MgGDP
MgGTP
MgGMPPCP
N = 1 (fixed)
Kd = 238 ± 12 µM
N = 1 (fixed)
Kd = 245 ± 48 µM
N = 1 (fixed)
Kd = 265 ± 33 µM

**S2 Fig. ITC isotherms with aminoglycosides**. Thermodynamics of aminoglycoside binding to APH(3')-IIb (**A**) WT or (**B**) M95L variant. The top panels show the differential heat released following baseline subtraction and the bottom panels show ITC binding curves.

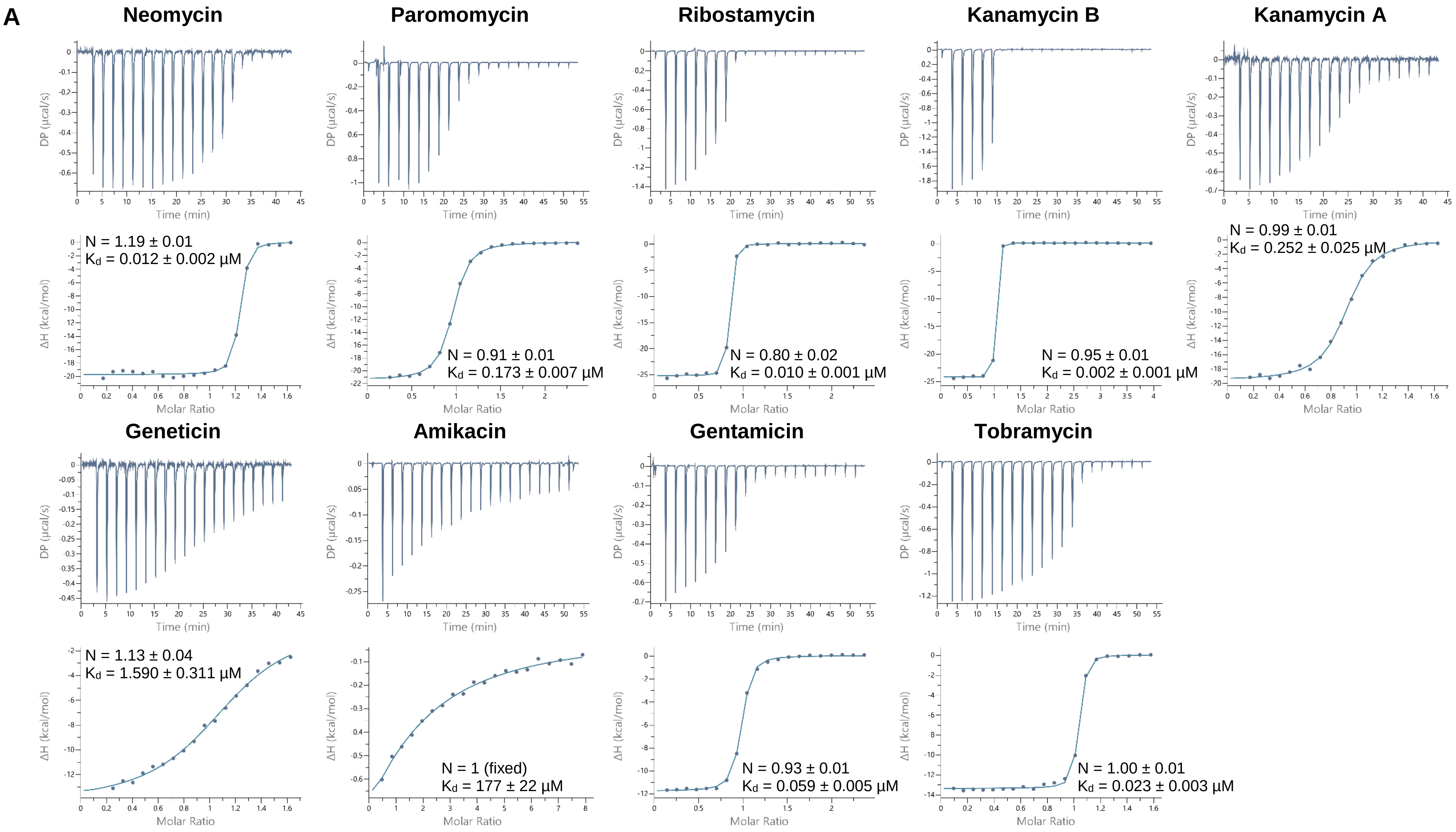

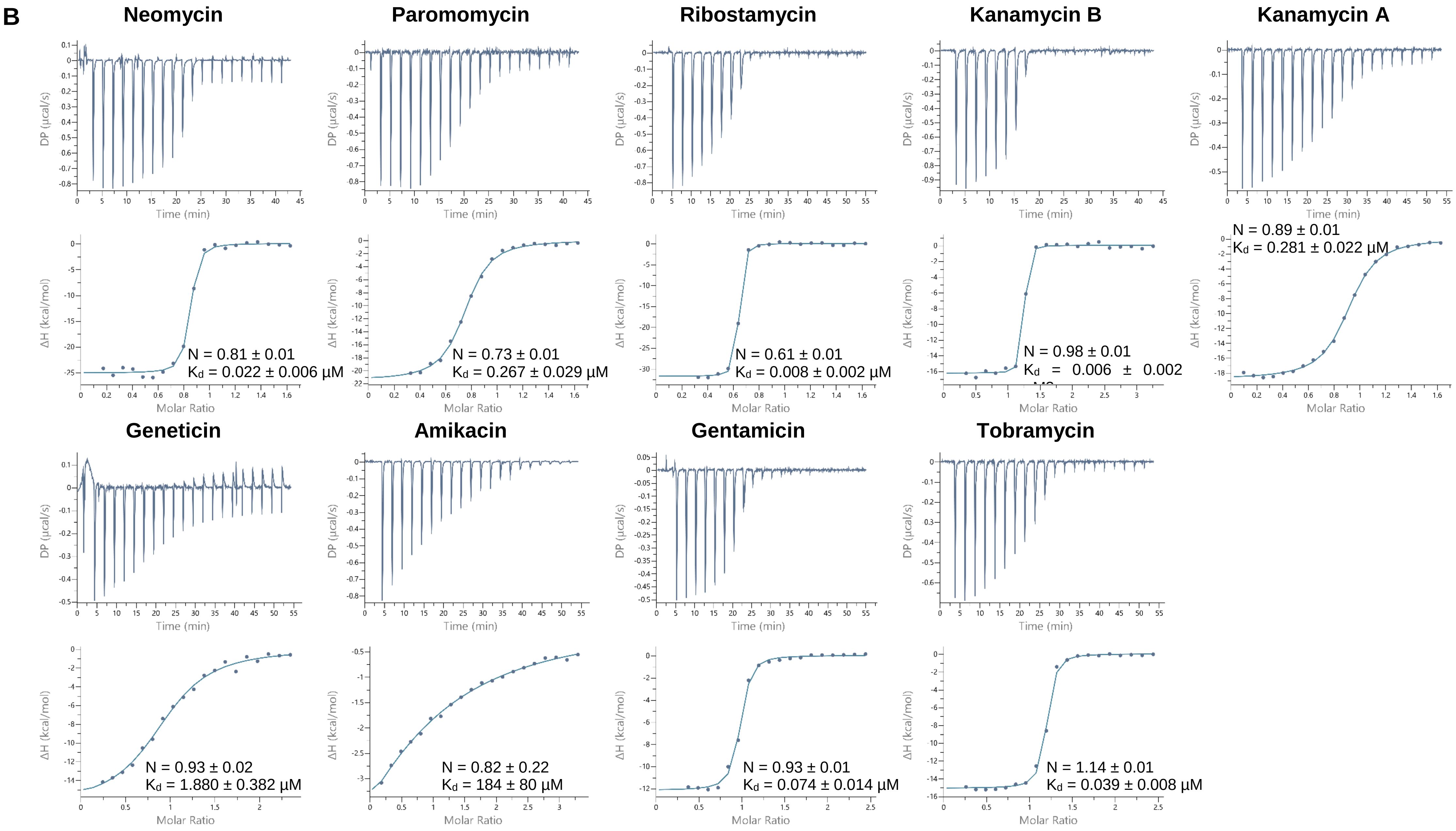
B
Neomycin
Paromomycin
Ribostamycin
Kanamycin B
Kanamycin A
DP (μcal/s)
Time (min)
ΔH (kcal/mol)
Molar Ratio
N = 0.81 ± 0.01
Kd = 0.022 ± 0.006 μM
N = 0.73 ± 0.01
Kd = 0.267 ± 0.029 μM
N = 0.61 ± 0.01
Kd = 0.008 ± 0.002 μM
N = 0.98 ± 0.01
Kd = 0.006 ± 0.002
N = 0.89 ± 0.01
Kd = 0.281 ± 0.022 μM
Geneticin
Amikacin
Gentamicin
Tobramycin
N = 0.93 ± 0.02
Kd = 1.880 ± 0.382 μM
N = 0.82 ± 0.22
Kd = 184 ± 80 μM
N = 0.93 ± 0.01
Kd = 0.074 ± 0.014 μM
N = 1.14 ± 0.01
Kd = 0.039 ± 0.008 μM

**S3 Fig. SAXS data**. Data were collected with 10 mg/mL of APH(3')-IIb in Tris buffer in absence of DTT (**A**) or in presence of 1 mM DTT (**B**). The two plots are overlayed in (**C**). The corresponding fitted data are shown in (**D**, **F**) and Guinier plots in (**E**, **G**).

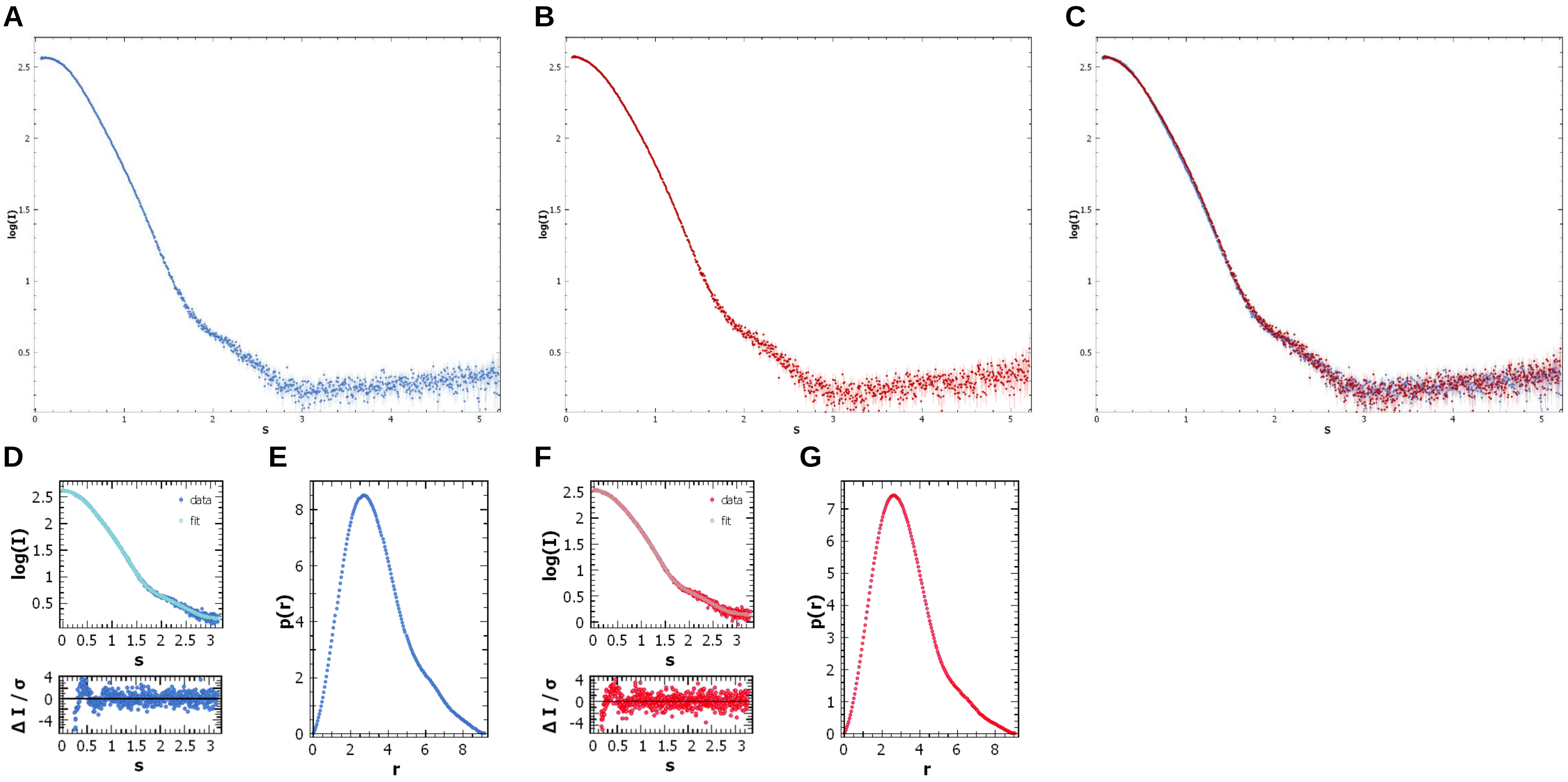

**S4 Fig. Comparison of crystal structures of different APHs**. (**A**) The high flexibility of the loop located between the β1 and β2 sheets is supported by the diversity of its conformation in APH(3')-IIa·Kanamycin (yellow, PDB 1ND4), APH(3')-Ic·ATP (purple, PDB 4EJ7) and APH(3')-IIIa·ADP·Kanamycin (blue, PDB 1L8T), and by the absence of electron density in APH(3')-IIb·ADP·KanaPhos (green, PDB in progress) and APH(3')-Id·ADP·Streptomycin (salmon, PDB 6FUX). Comparison of the conformational changes of the aminoglycoside binding loop in (**B**) APH(3')-IIb (apo in green and complexed with ADP and KanaP in pale green) and in (**C**) APH(3')-IIIa (apo in blue and complexed with ADP and Kana in pale blue). The aminoglycoside binding-induced shift of the Cα of (**B**) D162 or (**C**) E157 is indicated.

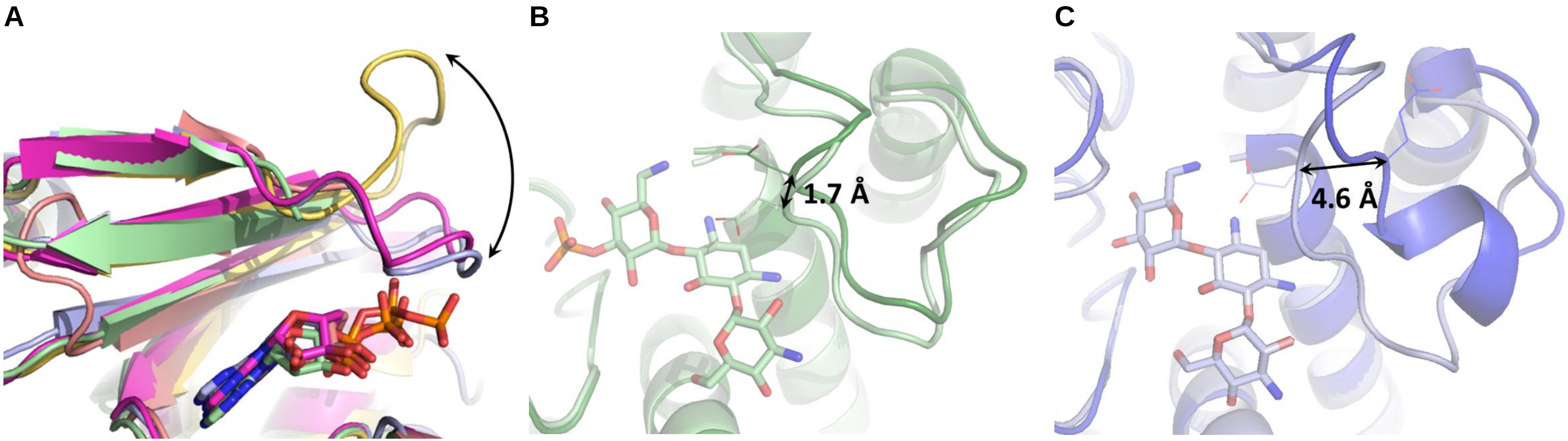

**S5 Fig. Prediction of interactions of APH(3')-IIb with 4,5- and 4,6-disubstituted aminoglycosides**. The ligands were placed using the electron density of PhosKanA using Coot. (**A**) PhosKanA template from the crystal structure. (**B**, **C**) Kanamycins A and B placed, with the yellow circle showing their difference. (**D**, **E**) geneticin from the crystal structure or placed, (**F**, **G**) neomycin and ribostamycin placed with the green frames showing potential additional interactions compared to kanamycin B, and blue circle the group that differs between neomycin and paromomycin. (**H**) Tobramycin placed with the red circle showing the missing 3'-hydroxyl group. (**I**) Gentamicin placed in reverse position with black circle showing the putative phosphorylated hydroxyl group instead of the missing 3'-hydroxyl group. (**J**) Amikacin placed with the purple frame showing clashes with the protein.

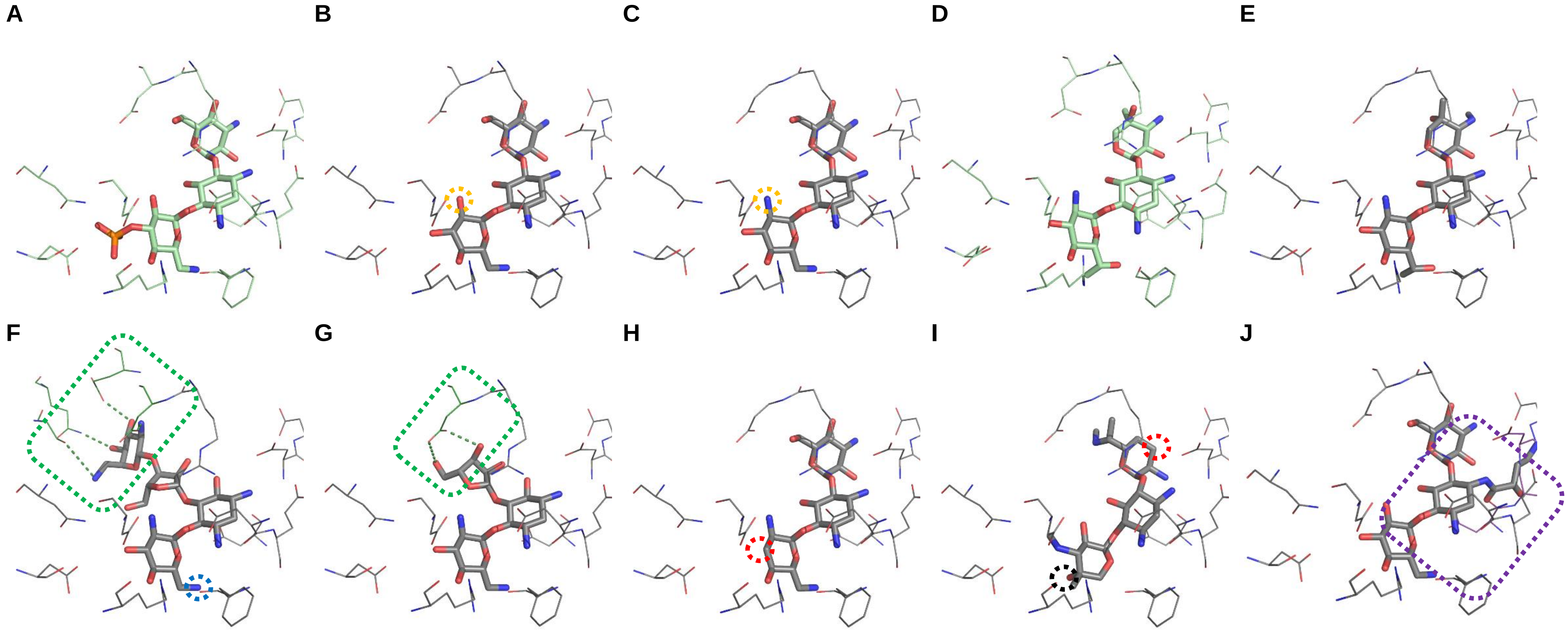
